# *Clostridioides difficile* binary toxin CDT induces biofilm-like persisting microcolonies

**DOI:** 10.1101/2024.05.23.595336

**Authors:** Jazmin Meza Torres, Jean-Yves Tinevez, Aline Crozouls, Héloïse Mary, Minhee Kim, Lise Hunault, Susan Chamorro-Rodriguez, Emilie Lejal, Pamela Altamirano-Silva, Samy Gobaa, Johann Peltier, Benoit Chassaing, Bruno Dupuy

## Abstract

Clinical symptoms of *Clostridioides difficile* infection (CDI) range from diarrhea to pseudomembranous colitis. A major challenge in managing CDI is the high rate of relapse. Several studies correlate production of CDT binary toxin by clinical strains of *Clostridioides difficile* with higher relapse rates. Although the mechanism of action of CDT on host cells is known, its exact contribution to CDI is still unclear. To understand the physiological role of CDT during CDI, we established two hypoxic relevant intestinal models, Transwell and Microfluidic Intestine-on-Chip systems. Both were challenged with the epidemic strain UK1 CDT^+^ and its isogenic CDT^-^ mutant. We report that CDT binary toxin induces mucin-associated microcolonies that increase *C. difficile* colonization and display biofilm-like properties by enhancing *C. difficile* resistance to vancomycin but not to fidaxomicin, a biofilm disrupting antibiotic. Importantly, biofilm-like CDT-dependent microcolonies were also observed in the caecum and colon of infected mice. Hence, our study shows that CDT toxin induces biofilm-like microcolonies, increasing *C. difficile* colonization and persistence.

## INTRODUCTION

*Clostridioides difficile* is a Gram-positive, obligate anaerobic and spore forming bacterium and the leading cause of antibiotic-associated diarrhea^1^. Antibiotic treatments altering the gut microbiota and reducing the production of secondary bile acids by the commensal microbiota allow germination of *C. difficile* spores present in the gut^2,3^. The clinical manifestations of *C. difficile* infection (CDI) range from diarrhea, to potentially fatal pseudomembranous colitis. Although CDI are mainly nosocomial, the incidence of community-associated CDI is rising^4,5^.

A major challenge in managing CDI is the high rate of relapse^6–8^. Recurrent CDI (rCDI) due to relapse or reinfection occur in 20-35% of cases in the two months following the initial episode^5,9,10^. After a first relapse episode, patients have a higher risk (around 60%) of presenting a second relapse^5,9,11^. Recent findings indicated that spores contribute to *C. difficile* persistence and rCDI^12^. *C. difficile* spores are able to entry into epithelial cells^12^, suggesting that they are liberated into the lumen upon epithelial cell renewal to potentially recolonize the host. However, inhibition of spore entry into epithelial cells only delayed relapse^13^, indicating that other mechanisms are involved in rCDI.

The key virulence factors of *C. difficile* involved in the host intestinal damages are the two large toxins TcdA and TcdB. These toxins are encoded on a pathogenicity locus (PaLoc) whose sequence variabilities define different *C. difficile* toxinotypes^14–18^. As monoglucosyltransferases, both toxins modify and inactivate Rho and Rac GTPAses^19^, triggering the disruption of the cytoskeleton, the breakdown of tight junctions and the subsequent loss of epithelial integrity^20^. In addition, 17-23% of clinical strains produce a third toxin, namely the *C. difficile* transferase toxin (CDT) or binary toxin. This toxin is composed of two separate toxin components: CDTa, the enzymatic ADP-ribosyl- transferase that depolymerizes F-actin, and CDTb, the cellular binding component that forms heptamers after proteolytic activation and translocates CDTa into the cytosol. The ADP ribosylation of actin by CDTa allows the formation of long microtubules protrusions that form a tentacle-like network on the surface of epithelial cells^21^. The actin depolymerization also leads to a misguided secretion of vesicles containing extracellular matrix (ECM) proteins such as fibronectin. Altogether, the microtubules protrusions and ECM-containing vesicles increase the adherence of *C. difficile* to epithelial cells^21,22^.

Despite the advances regarding the mechanism of action of CDT on host cells, the role of CDT during infection and disease remains unclear^23^. To date, CDT has been shown to enhance colonization^24^ and CDT^+^ strains correlate with an increased virulence leading to more severe diarrhea, increased pain, higher fatality rates and higher rCDI^25–31^. Several studies assessed the role of CDT during infection and colonization but all had limitations or bias, such as the use of (i) insertional CDT gene mutants with possible polar effects^31^, (ii) originally non-toxigenic TcdA^-^TcdB^-^CDT^+^ strains^32^, (iii) non-isogenic strains^33^, or (iv) a short infection time (2-3 days)^34^.

In this study, we used *C. difficile* epidemic strain UK1 and generated *tcdA*^-^*tcdB*^-^*cdtAB*^+/-^ isogenic mutant strains to elucidate the role of CDT during CDI. We show that *C. difficile* binary toxin CDT has a role in colonization through formation of 3D biofilm-like microcolonies structures in a 2D Transwell Intestinal model (TIM), a 3D Intestine on chip model (IoC) and in a mice infection model. These microcolony structures have biofilm-like properties such as increased resistance to antibiotics treatments. Our results support the implication of CDT in *C. difficile* long-term colonization and suggest that the 3D biofilm-like CDT-dependent structures are involved in *C. difficile* persistence in the gut. These findings provide evidence that CDT could play a crucial role in *C. difficile* relapses.

## RESULTS

### 1. CDT induces mucin-associated microcolonies

The CDT binary toxin triggers the formation of long microtubule protrusions and secretion of ECM-containing vesicles leading to increased *C. difficile* adherence^21,22^. The impact of CDT on epithelial cells has been studied by incubating the purified binary toxin with cell monolayers and *C. difficile* for short periods of time^22^. However, the impact this toxin could have on a longer period of time has not been explored. To tackle this question, we standardized two hypoxic cell culture models: a 2D Transwell Intestinal Model (TIM) with polarized cells under static conditions and a 3D Intestine-on Chip (IoC) microfluidic system that mimics flow and peristaltic intestinal motions. Both models were established with Caco-2 cells alone or Caco-2 cells co-cultured with HT29-MTX cells (mucus secreting cell-line) under hypoxia conditions (4% O2 and 5% CO2) optimized to simultaneously maintain viability of both eukaryotic and *C. difficile* cells (Fig. 1). Eukaryotic cell cytotoxicity, was monitored with lactate dehydrogenase (LDH) release assays and delta values between normoxia and hypoxia were calculated. Delta values ≤40% revealed that experiments of 24 or 48 h can be performed with the TIM and the IoC models, respectively, under these hypoxia conditions (Fig. S1A and S1B). Cell morphology, 3D or 2D structure and mucus production were assessed in these conditions, using appropriate markers (Fig 1B and 1D).

**Figure 1.**
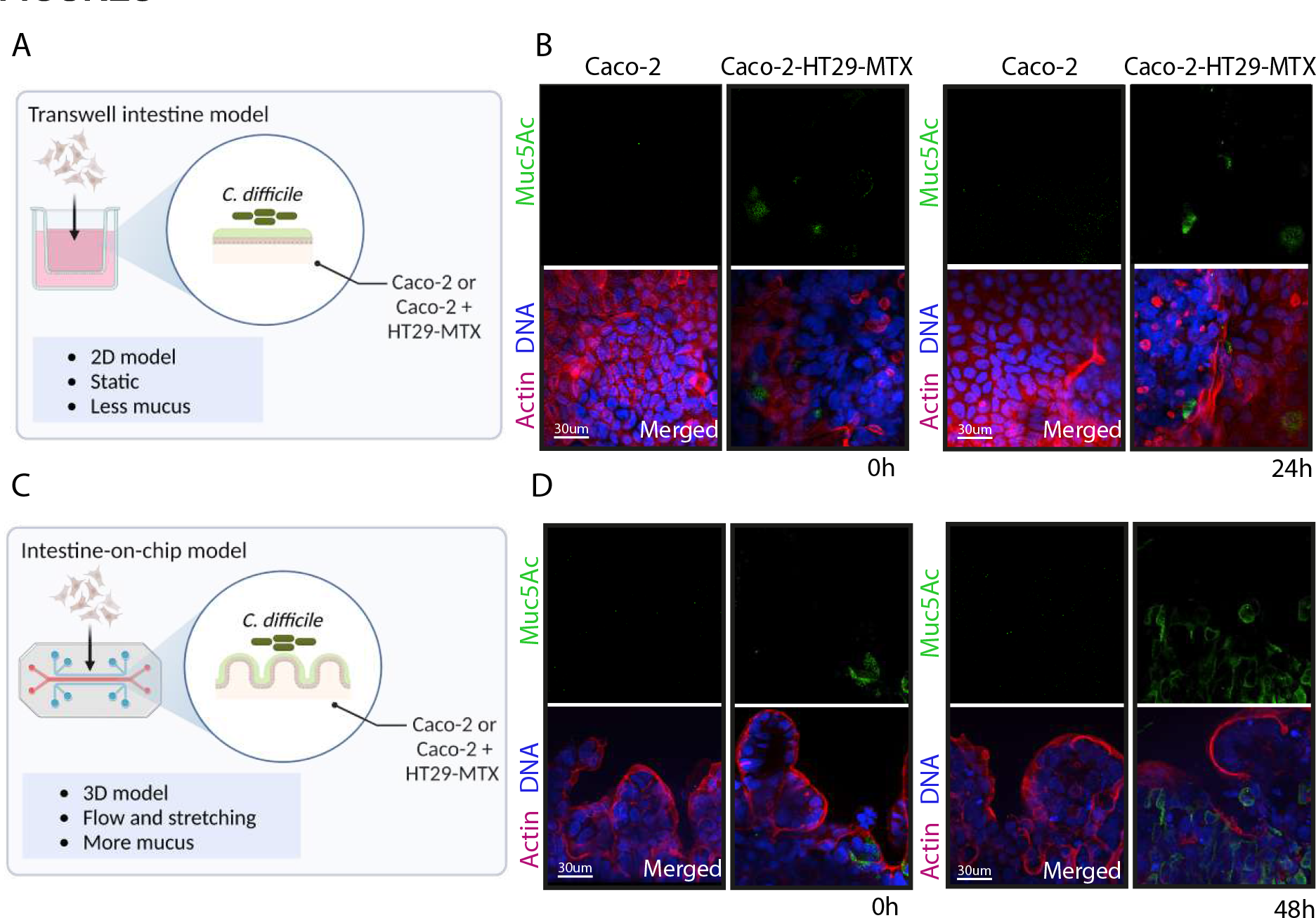
Establishment of hypoxic intestinal models to study the role of the CDT binary toxin during *C. difficile* infection. (A) Schematic representation of a Transwell Intestine Model (TIM) composed of Caco-2 cells alone or with HT29-MTX cells under hypoxia conditions (4% O2, 5% CO2). (B) Representative 3D reconstructed images of uninfected TIM under normoxia conditions (T0h, 5% CO2) and under hypoxia conditions (T24h, 4% O2, 5% CO2) (C). Schematic representation of the Intestine-on-chip model (IoC) composed of Caco-2 cells alone or with HT29-MTX cells under hypoxia conditions (4% O2, 5% CO2). (D) Representative 3D reconstructed images of uninfected IoC under normoxia conditions (T0h, 5% CO2) and under hypoxia conditions (T48h, 4% O2, 5% CO2). DNA was labelled with DAPI (blue), mucin with anti-Muc-5AC AF488 (green) and actin with phalloidin rhodamine (red).

To evaluate the role played by CDT in *C. difficile* colonization, two different strains were used: the reference strain 630Δ*erm* (TcdAB^+^CDT^-^) and the epidemic NAP1/B1/027 strain UK1 (TcdAB^+^CDT^+^). In order to assess CDT effects independently of TcdA and TcdB cytotoxic effects, we generated *in-frame tcdBEA* deletion mutants in 630 Δ*erm* (630 ToxAB^-^CDT^-^) and UK1 (UK1 ToxAB^-^CDT^+^) strains (FigS2A). Then, we generated an *in-frame cdtAB* deletion mutant in the UK1 ToxAB^-^ background (UK1 ToxAB^-^CDT^-^) (FigS2A). Deletion of either PaLoc or CdtLoc genes or both had no impact on *C. difficile* growth compared with the respective wild-type strains (Fig S2B). The absence of TcdA or CDT in culture supernatants of the ToxAB^-^ and CDT^-^ strains, respectively, confirmed the deletions (Fig S2C and S2D).

Caco-2 cells alone or co-cultured with HT29-MTX cells in the TIM model were first infected with *C. difficile* mutants (10^6^ bacteria/mL) and viable vegetative cells (CFU) and spores were numerated after 24h of infection (Fig S3A and S3B). No difference in CFU was observed between Caco-2 cells alone and Caco-2 cells co-cultured with HT29-MTX cells. The number of viable cells was similar between UK1 ToxAB^-^CDT^-^ and UK1 ToxAB^-^CDT^+^, with a 2-log increase at 24h, whereas no CFU increase was observed for 630 ToxAB^-^CDT^-^ (Fig S3A). Next, *C. difficile* adhesion was monitored at different time points (3, 6, 18 and 24h) in the TIM model. The 630 ToxAB^-^CDT^-^ strain adhered much less than the two UK1 mutant strains. In addition, the UK1 ToxAB^-^CDT^+^ strain showed a significant better adhesion than the CDT^-^ isogenic strain at 24h post-infection (p.i.) (Fig S3C). Immunofluorescence (IF) microscopy was then carried out to visualize *C. difficile* cells in the TIM model. An anti-SlpA monoclonal antibody was used to label *C. difficile*^35^ and mucin was labeled with an anti-Muc-5AC antibody (Fig 2A). Overall, more bacteria were observed on Caco-2 cells co-cultured with HT29-MTX cells than Caco-2 cells alone at 24 h p.i. indicating that the presence of mucin producer cells stimulates *C. difficile* adhesion (Fig 2A). This result is consistent with the previously reported close association of *C. difficile* with mucin^36,37^. Whereas only few 630 ToxAB^-^CDT^-^ and UK1 ToxAB^-^CDT^-^ were detected on Caco-2 cells co-cultured with the HT29-MTX, a high number of UK1 ToxAB^-^CDT^+^ bacteria, organized as microcolonies or clumps and colocalizing with Muc-5AC was observed (Fig 2A and Fig. S4A). By applying a quantitative approach, we determined that the number of bacteria increased up to 10 times in the presence of CDT and mucin producer cells (Fig 2B). The bacterial surface of these microcolonies associated to the coculture of Caco-2 and HT29-MTX cells ranged from 200 to 3000 µm^2^ (Fig 2C) and suggested a 3D biofilm-like structure. Consistently, mucin has recently been shown to induce formation of *C. difficile* biofilm^38^ and to chemoattract *C. difficile*^37^.

**Figure 2.**
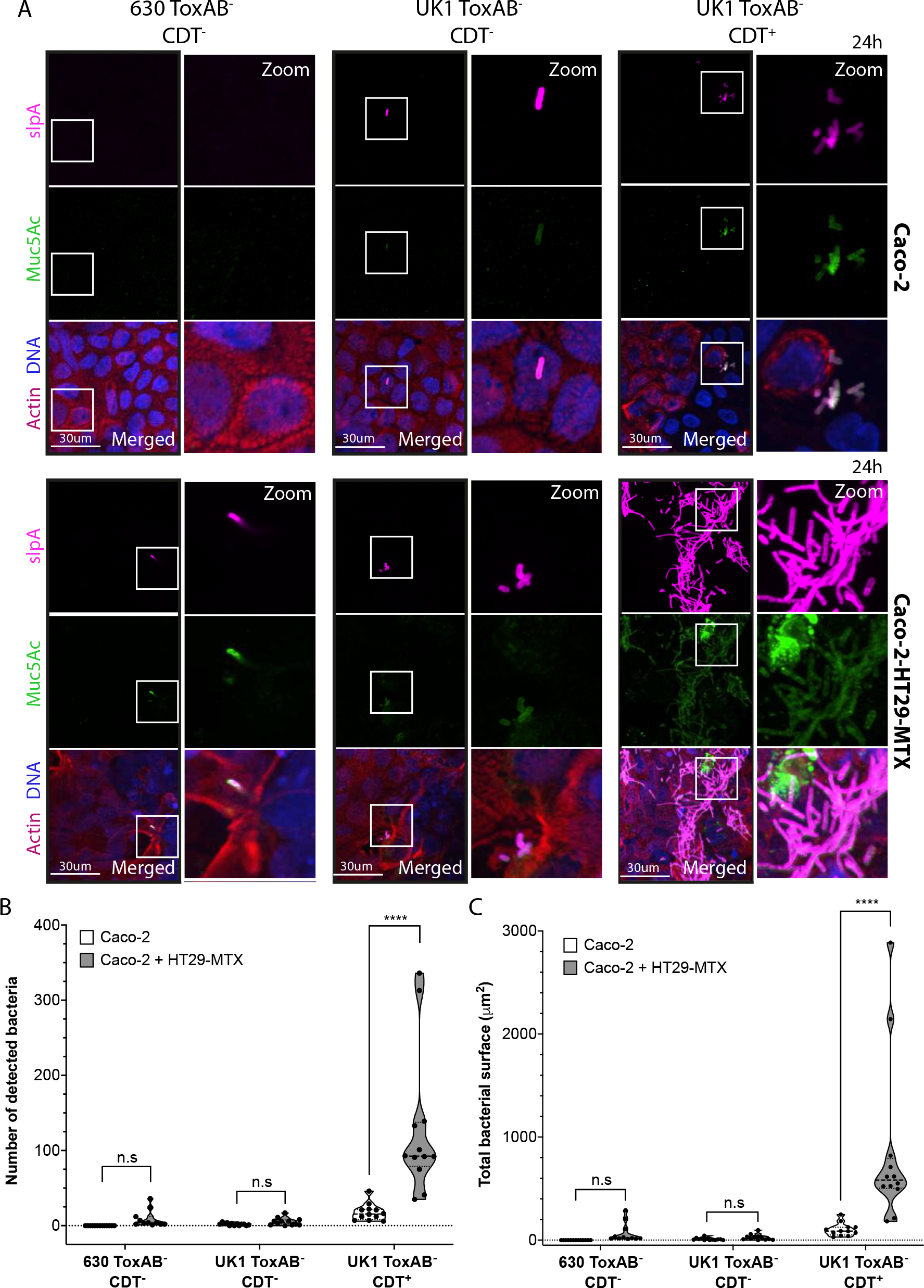
*C. difficile* forms CDT-mediated microcolonies in a Transwell Intestine model at 24h p.i. (A) Representative 3D reconstructed images of Caco-2 alone or with HT29-MTX cells infected with 630 ToxAB^-^CDT^-^, UK1 ToxAB^-^CDT^-^, or UK1 ToxAB^-^CDT^+^ during 24h under hypoxic conditions (4% O2, 5% CO2). DNA was labelled with DAPI (blue), mucin with anti-Muc- 5AC AF488 (green), actin with phalloidin rhodamine (red) and *C. difficile* with anti-SlpA^35^ AF647 (magenta). (B) Number of bacteria detected 24h p.i in Caco-2 cells alone or with HT29- MTX cells infected with different *C. difficile* strains as indicated. (C) Total bacteria surface detected 24h p.i in Caco-2 cells alone or with HT29-MTX cells infected with different *C. difficile* strains as indicated. The number of bacteria and total bacterial surface detected are reported for each image and at least 10 images were quantified per condition. Each black circle in the graph represents one image. Data and quantifications are representative of 3 independent biological replicates. Multiple unpaired *t* tests were performed and statistical significance is represented with **** (p<0.0001).

To further validate that mucin-associated microcolonies formation is specifically induced by the CDT toxin, we next performed a rescue experiment in the TIM model. For this, we infected Caco-2 cells co-cultured with HT29-MTX cells with UK1 ToxAB^-^ CDT^-^ or 630 ToxAB^-^CDT^-^ strains and exogenously supplied the purified CDT toxin^39^ at 6h p.i or at the time of the infection, corresponding to 18 and 24h treatment, respectively. Addition of purified CDT led to the formation of strong microcolonies for both CDT^-^ strains (Fig 3A and 3B). When CDT was added at 18h p.i, corresponding to 6h treatment, bacteria were not able to form microcolonies (Fig S5A and S5B). The number of detected bacteria (up to 300 bacteria per image) and the microcolony surface (up to 2000 µm^2^) observed after 18h of CDT treatment (Fig. 3C and 3D) were highly similar to those obtained with UK1 ToxAB^-^ CDT^+^ at 24h p.i. (Fig 2B and 2C). Cell exposure to CDT during 24h dramatically increased the number of adhered bacteria (up to 1000 bacteria per image) and the surface of the microcolonies (up to 6000 µm^2^ per image) (Fig. 3B, 3C and 3D). Formation of similar mucin-associated microcolonies by the strain 630 ToxAB^-^, naturally lacking the CDT toxin provides robust evidence that CDT is the sole factor involved in their formation. Altogether, these results strongly support that CDT toxin enhances close association of *C. difficile* with host cells, increases bacterial adhesion in presence of mucin and allows subsequent formation of microcolonies. These data thus suggest that the CDT-mediated microcolonies could contribute to *C. difficile* gut colonization.

**Figure 3.**
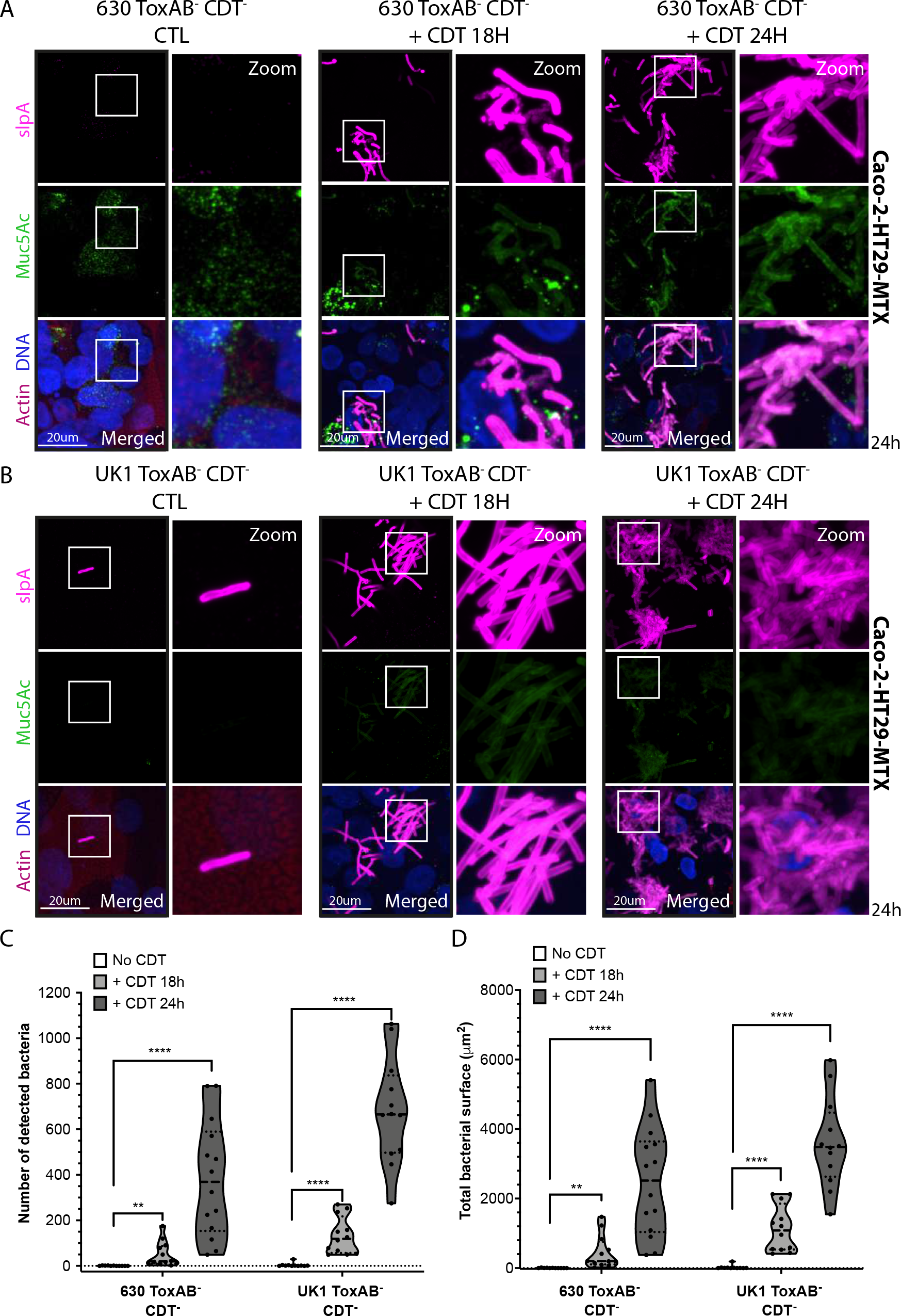
Purified CDT toxin induces microcolonies formation in CDT^-^ strains in the Transwell intestine model. Representative 3D reconstructed images of Caco-2 cocultured with HT29-MTX cells infected with (A) 630 ToxAB^-^CDT^-^ or (B) UK1 ToxAB^-^CDT^-^ during 24h under hypoxic conditions (4% O2, 5% CO2). Infected intestinal cells were exposed to CdtA (200ng/mL) and activated CdtB (400 ng/mL) during 18h or 24h. DNA was labelled with DAPI (blue), mucin with anti-Muc5AC AF488 (green), actin with phalloidin rhodamine (red) and *C. difficile* with anti-SlpA^35^ AF647 (magenta). (C) Number of bacteria detected 24h p.i in Caco-2 cells cocultured with HT29-MTX cells infected with *C. difficile* strains as indicated. (D) Total bacteria surface detected 24h p.i in cells infected with *C. difficile* strains as indicated. The number of bacteria and total bacterial surface detected are reported for each image and at least 10 images were quantified per condition. Each black circle in the graph represents one image. Data and quantifications are representative of 3 independent biological replicates. Multiple unpaired *t* tests were performed and statistical significance is represented with **(p≤0.01), **** (p≤0.0001).

### 2. CDT-dependent 3D microcolonies favor *C. difficile* colonization

To better decipher the role played by CDT in microcolony formation and *C. difficile* gut colonization, we next used the IoC model cultured with Caco-2 cells alone or in combination with HT29-MTX cells. Upon the development of the 3D intestinal structure (6-7 days after seeding in normoxic conditions), the IoC chips were placed in hypoxia (4% of O2) and infected with 630 ToxAB^-^CDT^-^, UK1 ToxAB^-^CDT^-^ or UK1 ToxAB^-^CDT^+^ for 24 to 48h. IF analyses revealed the formation of 3D microcolonies in the IoC model only with Caco-2 cells co-cultured with HT29-MTX cells at 24 and 48h p.i with the CDT^+^ strain but not with the CDT^-^ strains (Fig. 4A and 4B, Fig. S6A and S6B). CDT^+^ strains showed a higher number of detected bacteria and a higher bacterial surface when compared to the CDT^-^ strains (Fig. 4C and 4D, Fig S6C and S6D). The number of bacteria and surface of microcolonies observed with the IoC model at 48h. p.i., although slightly lower, were consistent with those obtained with the TIM model after 24h of infection (Fig 2B and 2C). Moreover, the microcolonies observed in IoC colocalized with mucin as in the TIM model (Fig S4A and S4B). The delay in microcolony formation in the IoC model when compared with the TIM can be explained by the presence of a flow. Nonetheless, the IoC model confirms that CDT promotes *C. difficile* colonization through the formation of 3D mucin-associated microcolonies.

**Figure 4.**
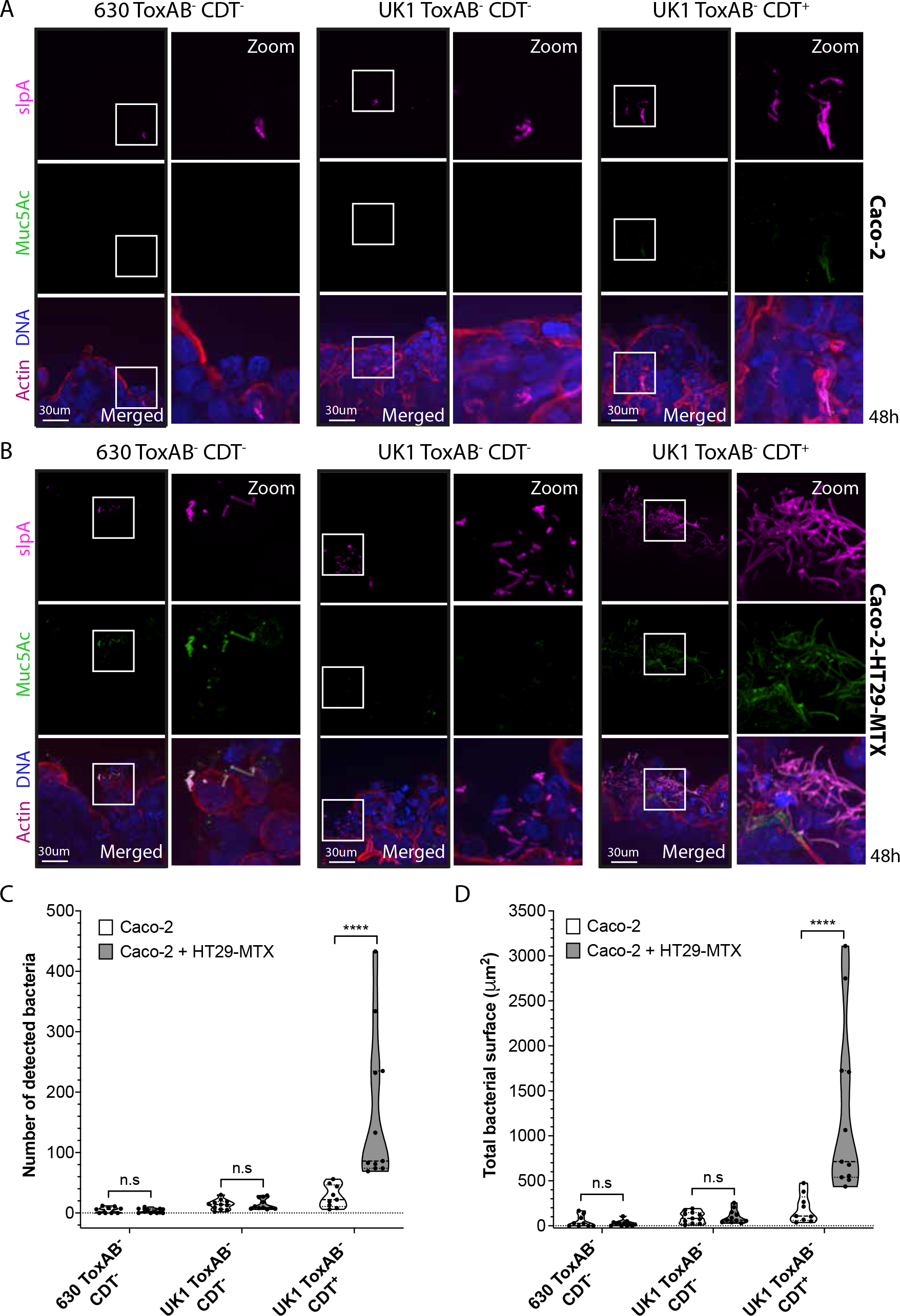
*C. difficile* forms CDT-mediated microcolonies in an Intestine-on-chip model at 48h p.i. (A) Representative 3D reconstructed images of Caco-2 cells (A) or Caco-2 cells cocultured with HT29-MTX cells (B) infected with 630 ToxAB^-^CDT^-^, UK1 ToxAB^-^CDT^-^ or UK1 ToxAB^-^CDT^+^ during 48h under hypoxic conditions (4% O2, 5% CO2). DNA was labelled with DAPI (blue), mucin with anti-Muc5AC AF488 (green), actin with phalloidin rhodamine (red) and *C. difficile* with anti-SlpA AF647 (magenta). (C) Number of bacteria detected 48h p.i in Caco- 2 cells alone or with HT29-MTX cells infected with *C. difficile* strains as indicated. (D) Total bacteria surface detected 48h p.i in Caco-2 cells alone or with HT29-MTX cells infected with *C. difficile* strains as indicated. The number of bacteria and total bacterial surface detected are reported for each image and at least 10 images were quantified per condition. Each black circle in the graph represents one image. Data and quantifications are representative of 2 independent biological replicates. Multiple unpaired *t* tests were performed and statistical significance is represented with **** (p<0.0001).

### 3. CDT-dependent microcolonies possess biofilm-like properties

Bacteria embedded in biofilms have increased resistance to antibiotics, antimicrobial peptides and oxidative stresses^40,41^. *In vitro*, *C. difficile* biofilms display higher survival than planktonic cells when exposed to antibiotics widely used to treat CDI, including vancomycin^42,43^. *C. difficile* biofilms can withstand vancomycin concentrations up to 25 times higher than the Minimal Inhibitory Concentration (MIC)^44^. On the other hand, fidaxomicin, an effective antibiotic against CDI and known to decrease rCDI, is effective in disrupting *C. difficile* biofilms^44,45^. Since CDT is a virulence factor associated with higher relapse rates^29^, we hypothesized that CDT-induced microcolonies could present biofilm-like properties, enabling a better resistance of *C. difficile* to vancomycin but not fidaxomicin. No difference in the MIC of fidaxomicin and vancomycin against the planktonic cells of UK1 ToxAB^+^CDT^+^, UK1 ToxAB^-^CDT^-^ and UK1 ToxAB^-^CDT^+^ strains grown in ADMEM medium was observed (Methods, Table S3), indicating that toxin gene deletions have no impact on resistance to these antibiotics. Caco-2 cells co-cultivated with HT29-MTX cells in the TIM model were then infected with UK1 ToxAB^-^CDT^-^ and UK1 ToxAB^-^CDT^+^ strains. After 24h infection, infected cells were treated with different concentrations of vancomycin or fidaxomicin (1x, 10x and 100x MIC) and incubated for an additional 24h before measuring viable CFU. No difference in CFU was observed between the two strains in absence of antibiotic treatment (CTL in Fig 5A and 5B). However, the CDT^+^ strain was significantly more resistant to vancomycin than the CDT^-^ strain for all concentrations tested (Fig 5A). Whereas no viable CDT^-^ bacteria was detected with the highest vancomycin concentration, CDT^+^ bacteria were still present at a concentration of 10^3^ CFU/mL (Fig 5A). In contrast, fidaxomicin similarly impacted the viability of the CDT^+^ and CDT^-^ strains with a strong reduction of CFU at 1x MIC and no bacteria were detected at 10 or 100x MIC concentrations (Fig 5B). Our result is explained by the fact that fidaxomicin is effective in eradicating *C. difficile* biofilms^44,45^. Altogether, these experiments show that CDT-induced microcolonies possess biofilm-like properties and resist to vancomycin but not fidaxomicin.

**Figure 5.**
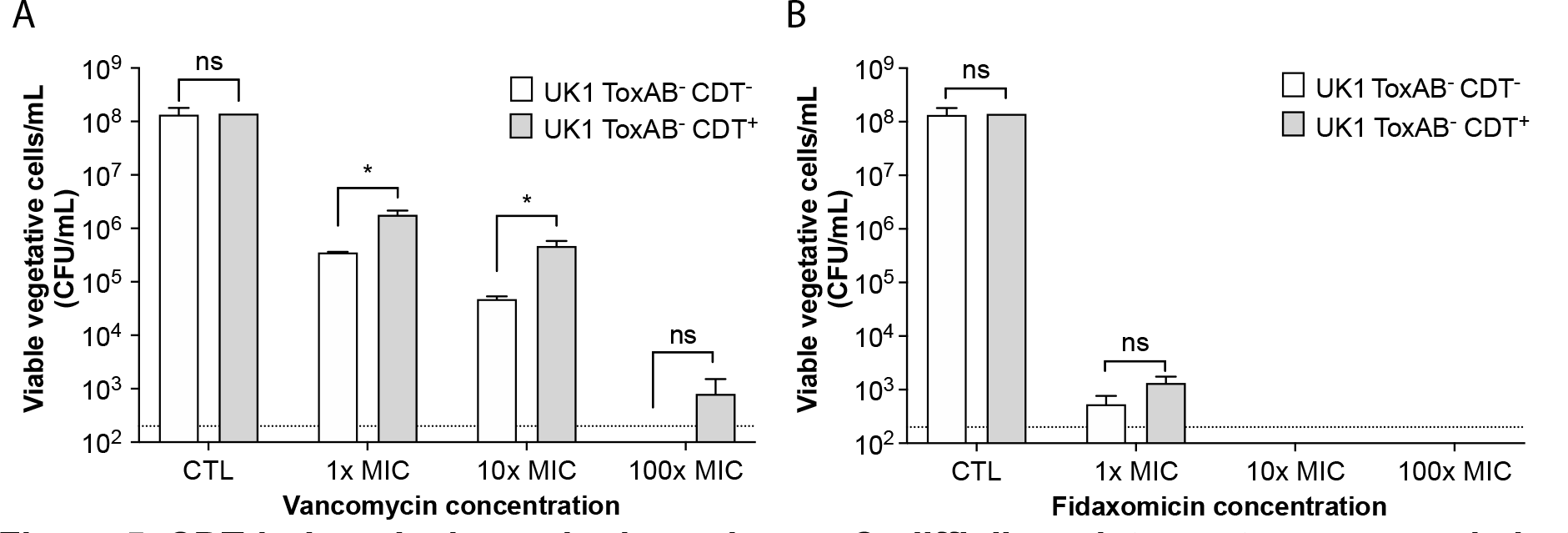
CDT-induced microcolonies enhance *C. difficile* resistance to vancomycin but not fidaxomicin. Caco-2 cocultured with HT29-MTX cells in the TIM model were infected with UK1 ToxAB^-^CDT^-^ and UK1 ToxAB^-^CDT^+^, 24h p.i infected cells were treated with different concentrations of vancomycin (A) or fidaxomicin (B) for additional 24h. Viable vegetative cells were recovered 48h p.i. The vancomycin and fidaxomicin concentrations used were 1x, 10x and 100x times higher than the MIC. Data represents mean with SEM from 3 independent biological replicates. Multiple unpaired *t* tests were performed and statistical significance is represented with * (p<0.05).

### 4. Mucin induces *C. difficile* biofilm formation *in vitro* and increases CDT levels

Induction of *C. difficile* biofilm by the presence of Muc2 in antibiotic-treated human fecal bioreactors has previously been reported by Engevik and collaborators^37^. Since biofilm formation in this study was evaluated with the CDT^+^ strain R20291, we next wondered whether Muc2-dependent biofilm formation *in-vitro* was mediated by CDT. To assess the role of CDT in biofilm formation, we cultured the strains 630 ToxAB^+^CDT^-^, UK1 ToxAB^+^CDT^+^, UK1 ToxAB^-^CDT^+^ and UK1 ToxAB^-^CDT^-^ during 48h in the Gut Microbiota Medium (GMM), a rich medium mimicking the intestinal milieu^46^, alone or with different types of mucins. Biofilm formation *in-vitro* was induced by the presence of native mucin and type II mucin in a CDT independent manner (Fig 6A). These data differ from the data obtained with the more physiologically relevant TIM and IoC models where the presence of both, CDT and mucin, was required to form biofilm-like microcolonies. Our result suggest that both mucin and CDT might induce *C. difficile* biofilm formation by different means.

**Figure 6.**
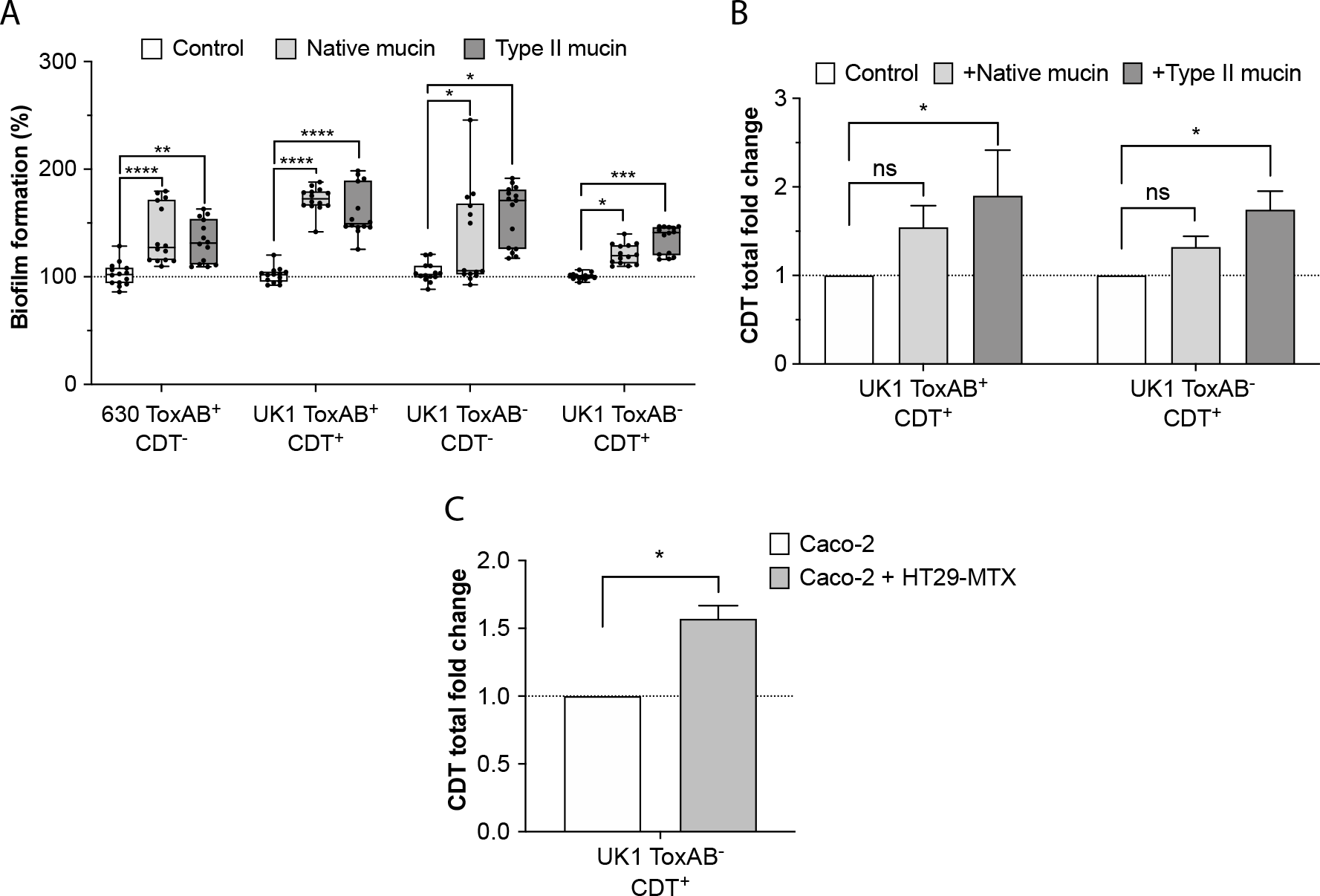
Mucin triggers *C. difficile* biofilm formation *in vitro* and increases the levels of CDT toxin. (A) Biofilm formation evaluated by crystal violet biofilm assay. Strains were grown during 48h in GMM medium alone (Control) or with native mucin from pork or mucin type II. The mean OD of biofilm from strains grown in GMM alone is adjusted to 100%. Minimum-maximum boxplot show 3 independent biological replicates with 14-15 technical replicates (black squares). Mann-Whitney U test was performed and statistical significance is represented (*p <0.05, ** p <0.01, ***p <0.001, ****p <0.0001). (B) CDT toxin ELISA represented as total CDT fold change. Strains were grown during 48h in GMM medium alone (Control) or GMM with native or mucin type II. The level of CDT assayed from crude extracts and supernatant were normalized to the OD600nm of bacteria cultures. Data represents mean with SEM from 3 independent experiments. A 2-Way ANOVA with Bonferroni correction was performed (ns: not statistically significant, *p <0.05). (C) CDT toxin ELISA represented as total CDT fold change. CDT was measured from supernatants recovered 24h p.i from Caco-2 alone or co-cultured with HT29-MTX cells in the TIM model infected with UK1 ToxAB^-^CDT^+^. Data represents mean with SEM from 2 independent experiments. An unpaired *t* test was performed and statistical significance is represented (* p<0.05).

We next wondered whether mucin could have an impact on CDT production. CDT levels were measured by enzyme-linked immunosorbent assay (ELISA) from supernatants and pellets collected after 48h of growth in GMM alone or with mucin. In both UK1 ToxAB^+^CDT^+^ and UK1ToxAB^-^CDT^+^ strains, CDT levels significantly increased in the presence of type II mucin (Fig 6B). Determination of the extracellular levels of CDT from supernatants of infected Caco-2 cells alone or co-cultured with HT29-MTX cells in the TIM model at 24h p.i. revealed a similar increase induced by the presence of mucin producer cells (Fig 6C). Thus, our data indicate that the presence of mucin increases CDT extracellular levels.

### 5. CDT toxin decreases mucin-related gene transcription

*C. difficile* has previously been shown to adhere to human mucus and to decrease mucin secretion in enteroids^47,48^. In addition, patients with CDI present decreased Muc2 levels and show alterations in mucin composition^48^. We therefore sought to determine whether the decreased mucin levels could be mediated by CDT. RNA were extracted from Caco-2 cells co-cultured with HT29-MTX cells treated with purified CDT (TIM and IoC models) and mucin mRNA levels were quantified by qRT-PCR (Fig 7). Cells from CDT-treated TIM model showed a significant decrease of *Muc2* and *Muc5AC* mRNA abundance genes after 6h but not 18h of CDT treatment compared to untreated cells (Fig 7A). The impact of the CDT treatment was delayed in the IoC model but a strong reduction of *Muc1*, *Muc2* and *Muc5AC* mRNA abundance was observed after 18h treatment (Fig 7B). The similar trend observed with both models indicates that CDT negatively regulates the mRNA abundance or stability of mucin-related genes. Altogether our results demonstrate that CDT has a double role in increasing both i) *C. difficile* adhesion to host cells leading to formation of 3D biofilm-like microcolonies and ii) the closeness to epithelial cells, the main target of *C. difficile* toxins, by reducing mucus production.

**Figure 7.**
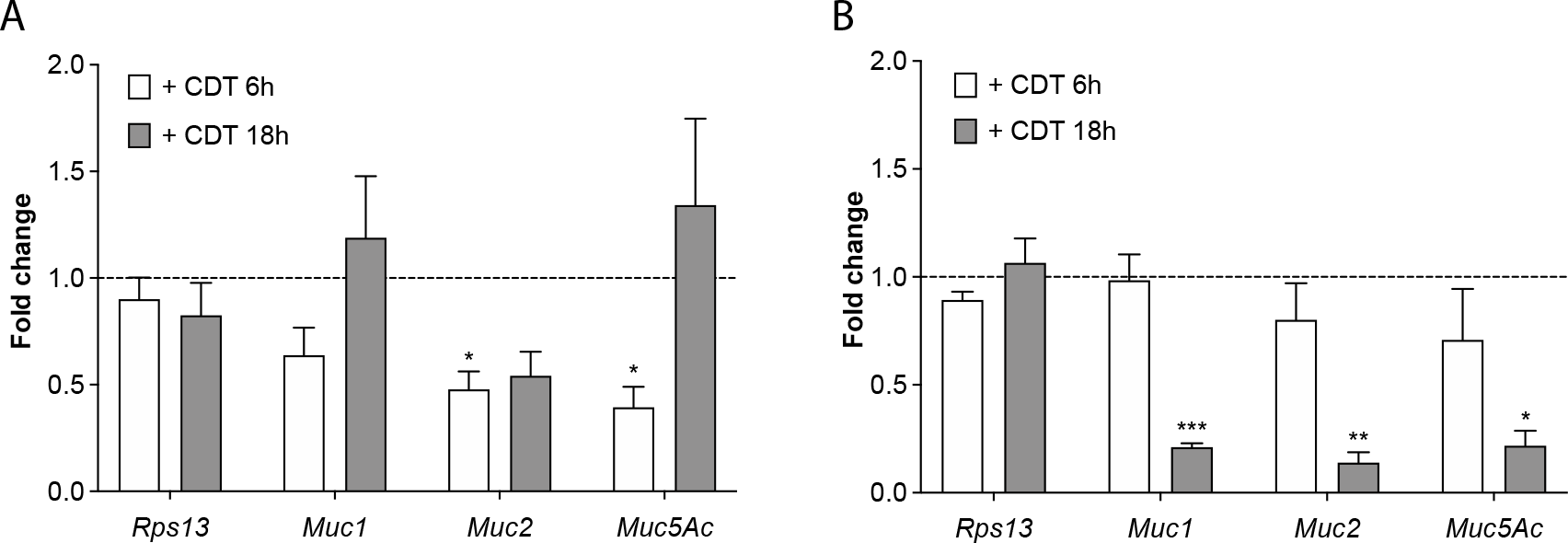
Cells treated with purified CDT reduce the transcription of mucin-associated genes. Caco-2 cells cocultured with HT29-MTX cells were treated or not with CdtA (200ng/mL) and activated CdtB (400 ng/mL), and mRNA levels of treated or not treated cells were quantified by qRT-PCR in (A) the Transwell intestine model (TIM) or (B) the Intestine-on-chip model (IoC). Data represents mean with SEM from 3 independent biological replicates (TIM) or 2 independent biological replicates (IoC) with 3 technical replicates. Data is normalized to not treated cells and represented as fold change relative to the housekeeping gene *Rps13*. A 2-way ANOVA with Geisser-Greenhouse correction test was performed and statistical significance is represented (* p<0.05, ** p <0.01 and, ***p <0.001).

### 6. CDT-infected mice harbor a decreased number of goblet cells, mucin thickness and present higher levels of inflammatory marker lipocalin-2

To determine whether CDT could influence *C. difficile* colonization and modify mucin levels *in-vivo*, C57Bl/6J germ-free mice (GFM) were infected with spores purified from the UK1 ToxAB^+^CDT^+^, UK1 ToxAB^-^CDT^-^ or UK1 ToxAB^-^CDT^+^ strains. Mice infected with UK1 ToxAB^+^CDT^+^ were sacrificed 2 days p.i due to their rapid loss of weight of around 25% (Fig S7A and S7B), as typically observed^49^. The number of total cells detected in the feces of mice, corresponding to the sum of vegetative cells plus spores, was similar for CDT^+^ and CDT^-^ strains during CDI (Fig8A). Remarkably, on day 6 p.i, the CDT^+^ strain showed significantly higher number of total cells than the CDT^-^ strain (Fig8A). Accordingly, when comparing only the vegetative cells, the CDT^-^ strain showed a decrease in CFU at day 6 p.i (Fig S7C). Inversely, a higher number of spores was recovered from the feces of mice infected with the CDT^-^ than with the CDT^+^ strain at days 6 and 7 p.i (Fig S7D). At day 8 p.i, mice infected with the CDT^-^ strain but not with the CDT^+^ strain completely cleared the vegetative bacteria (Fig S7C and S7D). Altogether, these results showed no significant difference in the number of vegetative cells released in the feces of mice between CDT^+^ and CDT^-^ strains (Fig 8A, S7C and S7D). However, CDT^+^ infected mice showed a delayed in clearing CDI compared to CDT^-^ strain (Fig S7C). In addition, significant differences were observed in the caecal content of mice sacrificed 13 days p.i, with a greater number of spores for the strain CDT^+^ (Fig S7E), supporting a better persistence of the CDT^+^ strain in the caecum of mice.

**Figure 8.**
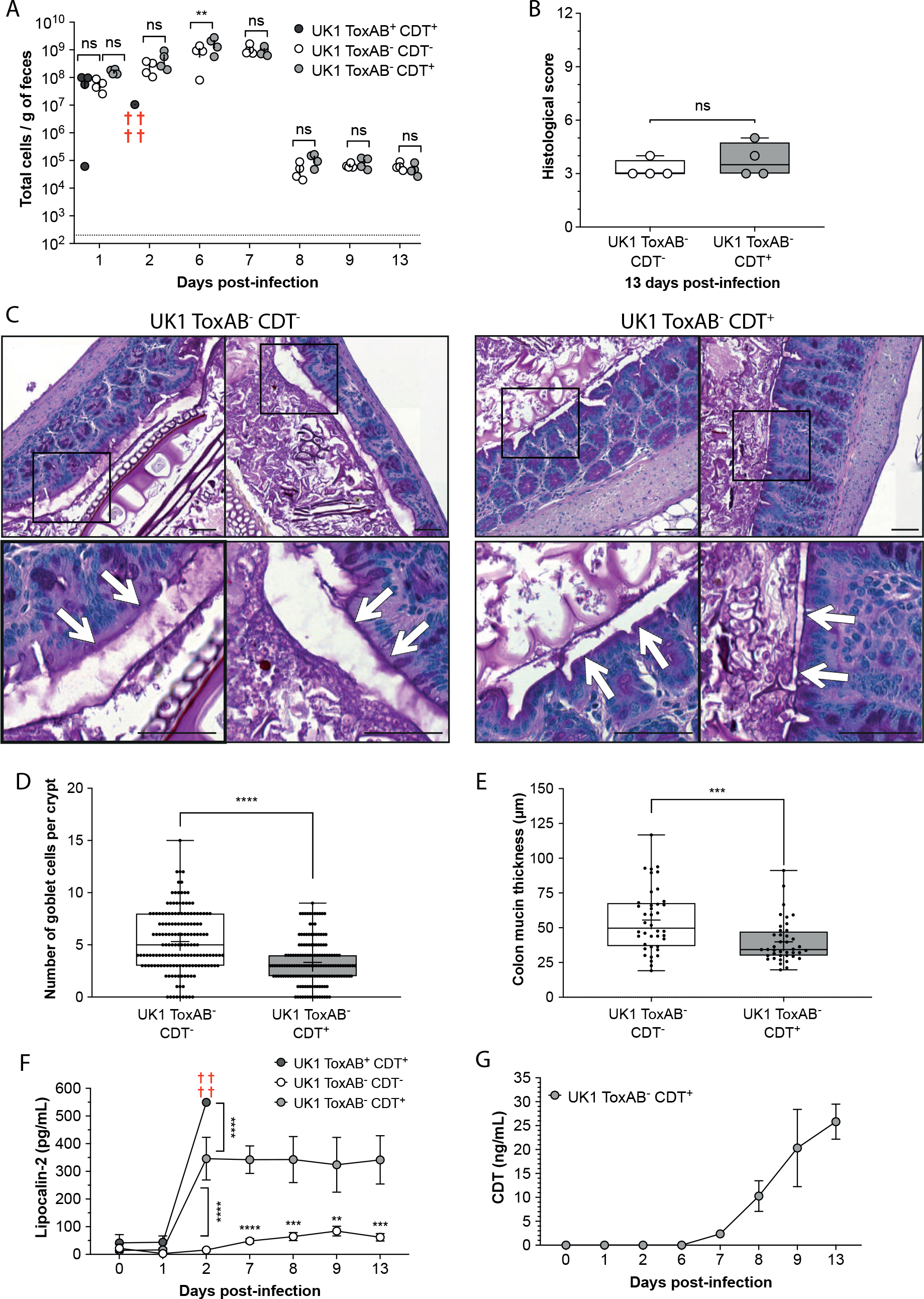
The CDT toxin increases gut inflammation and decreases mucin thickness and goblet cells numbers. (A) C57Bl/6J Germ-free mice were infected with purified spores from UK1 ToxAB^+^CDT^+^, UK1 ToxAB^-^CDT^-^ or UK1 ToxAB^-^CDT^+^ and the total number of CFUs (vegetative cells + spores) in feces was assessed at different days p.i. (B) Histological score 13 days p.i. (C) Representative images of mucin thickness (black bars) and goblet cells (dark purple) from mouse colonic sections 13 days p.i stained with PAS. (D) Quantification of number of goblet cells per crypt from mouse colonic sections 13 days p.i. (E) Quantification of mucin thickness from mouse colonic sections 13 days p.i. (F) Fecal lipocaline-2 levels detected by ELISA before (day 0 and 1) and during *C. difficile* infection (day 2, 6, 7, 8, 9 and 13). (G) Fecal CDT levels detected by ELISA before infection (day 0 and 1) and during *C. difficile* infection (day 2, 6, 7, 8, 9 and 13). Data represents mean with SEM (F, G). Multiple unpaired *t* tests (A, B, F) or Mann Whitney tests were performed (D, E) and statistical significance is represented with * p<0.05, ** p <0.01, ***p <0.001 and, **** p<0.0001. ns: no statistical significance.

No significant difference was found in intestinal inflammation, assessed by the histological score, between the CDT^+^ and CDT^-^ strains (Fig 8B). Colon and caecum were recovered and fixed with Carnoy to preserve mucin structure. Periodic Acid Schiff (PAS) staining in colon sections revealed a significant decrease in the number of goblet cells per crypt as well as mucin thickness in mice infected with the CDT^+^ strain compared to those infected with the CDT^-^ strain (Fig8C-E). This result further supports that CDT induces changes in mucin.

Toxin-induced inflammation is beneficial to *C. difficile* during infection^50^. In order to better understand the role of CDT in the toxin-dependent intestinal inflammation, we monitored the fecal lipocaline-2 (Lcn2) levels of mice infected with CDT^+^ and CDT^-^ strains. As expected, the strain ToxAB^+^CDT^+^ induced a huge inflammatory response 2 days p.i, whereas UK1 ToxAB^-^CDT^+^ and UK1 ToxAB^-^CDT^-^ strains showed intermediate or low levels of Lcn2, respectively (Fig 8F). The maximum inflammation occurred from day 2 p.i with a significant contribution of CDT and sustained up to day 13 when mice were sacrificed. Additionally, CDT levels monitored from mice feces increased from 7 up to 13 days p.i, suggesting that CDT levels are persistent during CDI (Fig 8G). Altogether these results show that CDT mediates changes in mucin thickness, decreases the number of goblet cells and induces an inflammatory response maintained throughout CDI.

### 7. CDT toxin favors the formation of biofilm-like microcolonies in the caecum and colon of mice, increasing *C. difficile* persistence

In order to study whether CDT biofilm-like microcolonies were also formed *in vivo,* mice colonic sections and caecum sections were immuno-stained 13 days p.i. Interestingly, mucin-associated and embedded microcolonies were detected in the caecum and colon of mice infected with *C. difficile* (Fig 9A and 9B). Moreover, the total surface of these microcolonies in the colon and caecum was significantly higher in mice infected with CDT^+^ than in those infected with CDT^-^, suggesting that CDT induces a better colonization in colon and caecum (Fig 9C). The size of the CDT-microcolonies was similar in the caecum and colon, from ∼100 µm^2^ up to ∼7000 µm^2^ per image, with a mean around 1000 µm^2^ (Fig 9C). The presence of CDT-associated microcolonies in both the colon and caecum epithelium of mice underscores their biological importance and suggests that they might be involved not only in *C. difficile* colonization but also in *C. difficile* persistence.

**Figure 9.**
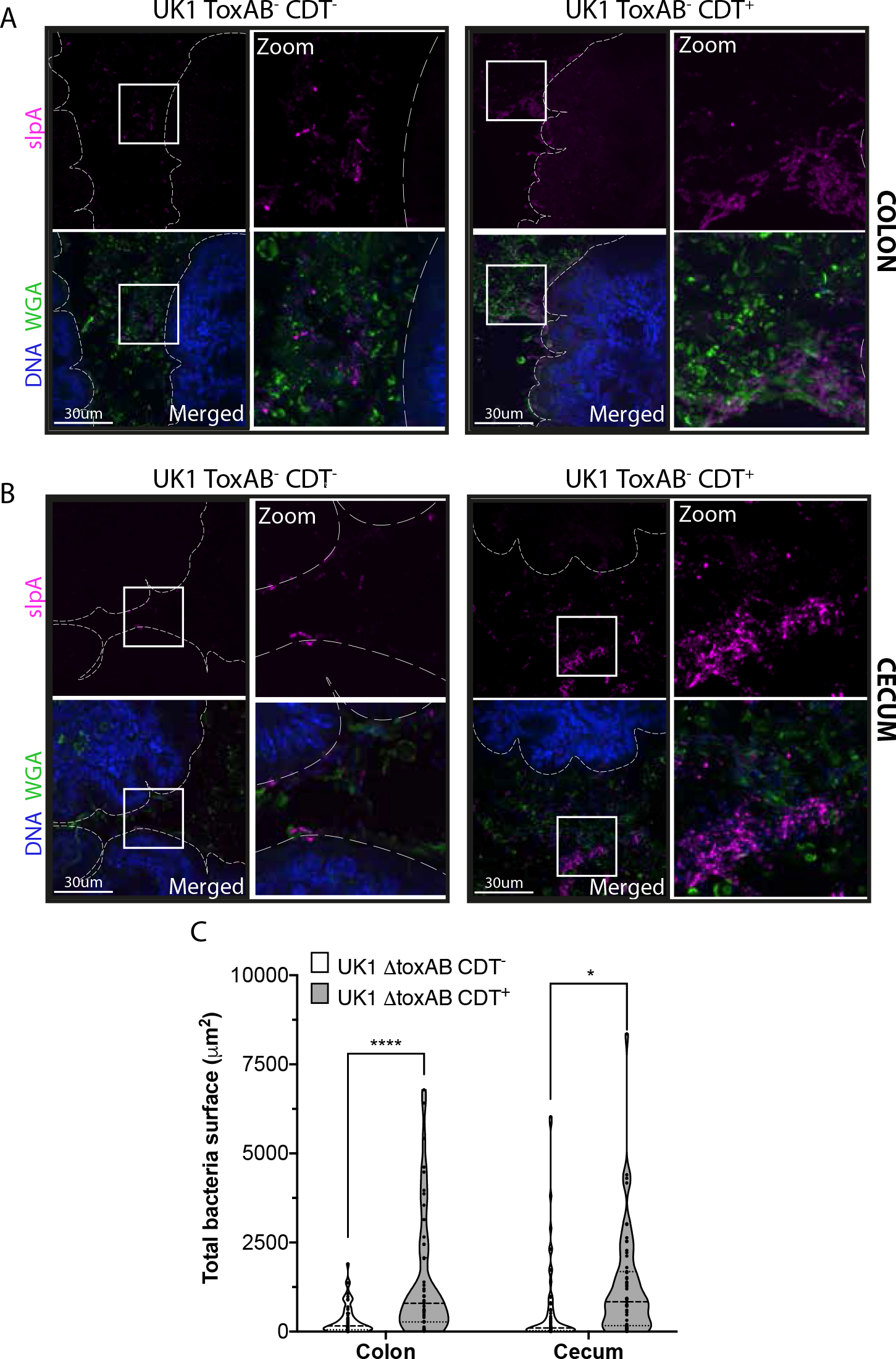
The CDT toxin promotes the formation of biofilm-like microcolonies in the caecum and colon of mice. (A) Mouse colonic and (B) caecum sections were immunostained 13 days p.i. DNA was labelled with DAPI (blue), mucus layer with the lectin WGA AF488 (green), and *C. difficile* was labelled with anti-slpA AF647 (magenta). (C) Total bacterial surface per image was quantified from mice colonic and caecum sections 13 days p.i. Data and quantifications are representative of at least 13 images quantified per mice (four mice per condition, scale bar 30 μm). Each black square in the graph represents one image (n ≥52). Multiple unpaired *t* tests were performed and statistical significance is represented (* p<0.05, **** p<0.0001).

## DISCUSSION

We report here that purified or secreted *C. difficile* CDT binary toxin induces mucin-associated microcolonies *in vitro*. CDT-induced microcolonies enhance *C. difficile* resistance to vancomycin but not to fidaxomicin, consistent with biofilm structures. Biofilm-like microcolonies are also formed *in vivo* in the caecum and colon of mice, facilitating *C. difficile* colonization and potentially promoting *C. difficile* persistence in mice. In addition, we showed that the presence of mucin increases CDT levels and that CDT induces, in turn, transcriptional changes of mucin**-**related genes resulting in a reduction of mucin thickness and goblet cells in the colon. Overall, we identified a new role of CDT during CDI that paves the way toward a better understanding of the association between CDT, the increased severity of CDI^30^ and the elevated recurrence rates of CDT^+^ strains ^29,51^.

The binary toxin CDT belongs to the family of binary actin-ADP-ribosylating toxins comprising toxins of other toxigenic Clostridia species^52–54^ responsible of gastrointestinal diseases in humans and/or animals^55^. All binary toxins share the same target i.e., the actin cytoskeleton^55^, a recurrent target of bacterial toxins^56^ that allow bacteria to create a replication niche^57^, prevent or induce phagocytosis or escape from the host immune system^58–60^. It has been shown that the binary toxin CDT promotes F-actin depolymerization and the formation of microtubules protrusions on epithelial cells thus increasing *C. difficile* adhesion to host cells. However, these studies were performed either with purified CDT in the absence of *C. difficile* or with cells pre-treated with CDT before infection with *C. difficile* for a brief incubation time (2h or 4h)^21,22,61^. In addition, these studies used polarized Caco-2 cells, which lack a mucus layer ^62^. On the contrary, our study included polarized Caco-2 cells co-cultured with HT29-MTX mucus producer cells, infected with a *C. difficile* CDT^+^ or CDT^-^ strain incubated with purified CDT under hypoxic conditions. Moreover, we used two models that better reflect the intestinal architecture. These experimental conditions revealed for the first time that CDT induced microcolonies with biofilm-like structure. We speculate that CDT activity, by inducing local modifications at the infection site, including on mucin and extracellular matrix, promote *C. difficile* attachment and cell growth, a prerequisite for *C. difficile* biofilm formation.

Biofilms are associated with persistent infections^63^ due in part to the capacity of bacteria embedded inside biofilms to be more resistant to antibiotics^40^ and host immune responses^64^. Thus, *C. difficile* strain R20291 is 10 times more resistant in biofilm than planktonic cells to vancomycin treatment^42^ and *C. difficile* clinical isolates are 100 times more tolerant to metronidazole^43,65^. *C. difficile* biofilms also increase resistance to oxygen levels, bile salts and antimicrobial peptides^43,66^. *C. difficile* can be found in multi-species biofilms formed by the gut microbiota, constituting a potential reservoir leading to asymptomatic carriage and risk of recurrent infection after antibiotic therapy^43,65,67–69^. Whether formation of a *C. difficile* mono-species biofilm can trigger a multispecies biofilm remains an open question. However, we know that formation of *C. difficile* mono-species biofilm occurs *in vitro*^68,70^ and is induced in response to sub-inhibitory concentrations of antibiotics or metabolites whose concentrations vary during gut dysbiosis^43,70,71^. We showed that the binary toxin CDT liberated during biofilm formation regulates the expression of mucin-related genes. Since toxins A and B are produced inside *C. difficile* biofilms^72^, CDT might have a dual impact by i) triggering biofilm formation and ii) acting as a virulence factor by making the gut epithelium more accessible to toxins A and B, thus increasing CDI severity^27,30^.

The release of mucin and antimicrobial molecules is regulated by the commensal microbiota^73^. Pathogens can also induce changes in secretion of mucin by goblet cells^74–76^, glycosylation of mucins^77^ or reduction in mucus viscosity^78^. Several studies have shown that mucin facilitates *C. difficile* colonization ^37,38,47,48,79^, probably because *C. difficile* binds to mucus from mice and humans^37,47,67,80,81^. Engevik *et al.* observed that *C. difficile* strain BAA-1878 can decrease Muc2 secretion when injected in human intestinal organoids (HIOs)^48^. They also showed a relationship between patients with recurrent CDI and a decrease of Muc2 and *N*-acetylgalactosamine (GalNAc) expression, along with an increase of *N*-acetylglucosamine (GlcNAc) and galactose residues^48^. Nonetheless, this study did not associate these changes in mucin with the capacity of the *C. difficile* strain BAA-1878 to produce CDT^82^. Our data provide evidence that CDT alone can directly or indirectly alter the level of intestinal mucus by a mechanism that remains to be investigated. We showed a transcriptional drop of mucin-associated genes in the TIM and IoC models in presence of CDT (Fig 7), and a decrease in mucin thickness and goblet cells in a murine model when infected with the UK1 ToxAB^-^CDT^+^ strain (Fig 8). Decreasing mucin transcription through CDT could be one of the mechanisms used by *C. difficile* to reach the gut epithelium. It is known that *in vivo C. difficile* uses in priority Stickland-acceptor amino-acids such as serine, proline and threonine ^49,83^. Recently, Furtado *et al.,* showed that uptake of serine and threonine is upregulated in *C. difficile* in presence of mucus^38^. Both of them are the main amino-acids of the mucin peptide backbone^84^. Intriguingly, lack of threonine resulted in a decrease of the mucus layer and goblet producer cells^85^. Therefore, we can speculate that mucus-associated microcolonies of *C. difficile* result to a decrease of the threonine levels leading to a reduction of the mucus layer thickness and shrinkage of goblet cells^86^.

Our data also showed that CDT expression is upregulated in the presence of mucin or mucin sugar derivates. However, the question of which specific mucin-derived monosaccharide(s) or polysaccharides(s) induce CDT expression remains open. In agreement with our data, CDT levels in patients are 20-fold higher than the CDT levels measured *in vitro*^87^. This finding is not unprecedented since expression of the cytolethal distending toxin (*cdtABC*) and vacuolating cytotoxin from *Campylobacter jejuni*^88^ are also upregulated in response to mucin. Interestingly, *C. jejuni* shows the same chemiotaxis towards mucin^89^, as recently demonstrated for *C. difficile* ^79^. Furthermore, toxins secreted by many pathogens can diffuse through the mucus, leading to the reduction of its production and subsequently, to the disruption of the epithelial barrier and intracellular tight junctions. These damages can compromise the mucosal barrier and promote invasion by pathogens ^86^. The binary toxin CDT might be strategically used by *C. difficile* to establish a biofilm embedded inside the mucus layer and surrounded by microtubule protrusions. Such biofilm could promote *C. difficile* persistence into the host by increasing cell surface adhesion and aggregation to resist shear forces and flow^90^, by allowing nutrient perfusion and providing protection against antimicrobial agents^91,92^, high oxygen tensions^93^, bile salts, and oxygen radicals ^94,95^. Within the biofilm, the CDT-mediated alteration of mucus, allows a closer proximity of the main toxins with the epithelium cells, which probably participates in the disease severity of patients infected by CDT^+^ strains of *C. difficile*.

Overall, our study provides new insights on the role of the binary toxin CDT during CDI, supporting previous studies that correlated the production of this toxin by *C. difficile* clinical strains with higher virulence and rCDI^25–31^. The unexpected role of CDT toxin in formation of biofilm-like mucin-associated microcolonies, opens new perspectives regarding the methods used by enteric pathogens to create a niche in the gut epithelium as a way to persist.

## METHODS

### Bacterial strains and culture conditions

Bacterial strains and plasmids used in this study are listed in table S1. *C. difficile* strains were routinely cultured on BHI agar (Difco), BHI broth (Difco), or TY broth (Bacto tryptone 30 g.L^-1^, yeast extract 20 g.L^-1^, pH 7.4) at 37°C in an anaerobic environment (90% [vol/vol] N2, 5% [vol/vol] CO2, and 5% [vol/vol] H2). When necessary, *C. difficile* culture media were supplemented with cefoxitin (Cfx; 25 mg/liter), cycloserine (Ccs; 250 mg/liter), thiamphenicol (Tm; 7.5 mg/liter), and erythromycin (Erm; 5 mg/liter). *Escherichia coli* strains were cultured at 37°C in LB broth or LB agar (MP Biomedicals), containing chloramphenicol (25 mg/liter), and when needed, ampicillin (100 mg/liter).

### Construction of *C. difficile* mutant strains

All primers used in this study are listed in table S2. A pathogenicity locus (PaLoc)- deleted strains of *C. difficile* 630Δ*erm* and UK1 that lacked the *tcdB*, *tcdE* and *tcdA* genes were generated^96^. A CDT locus (CdtLoc)-deleted strain UK1 strain, lacking the *cdtA* and *cdtB* genes (designed as CDT^-^), was then generated in the Δ*toxAB* background. The deletion mutants were created using a toxin-mediated allele exchange method^97^. Briefly, approximately 850 bp of DNA flanking the region to be deleted were amplified by PCR from *C. difficile* UK1 and 630Δ*erm*. Purified PCR products were cloned into the PmeI site of the pMSR0 vector using NEBuilder HiFi DNA Assembly (New England Biolabs). The resulting plasmid was transformed into *E. coli* strain NEB10β (New England Biolabs) and insert verified by sequencing. Plasmids were then transformed into *E. coli* HB101(RP4) and transferred by conjugation into the appropriate C*. difficile* strains. Transconjugants were selected on BHI supplemented with cycloserine, cefoxitin, and thiamphenicol. Allelic exchange was performed as described previously^97^.

### Cell culture

Caco-2 cells (clone TC-7) and HT29-MTX cells were provided by Nathalie Sauvonnet from Institut Pasteur, Paris, France. Cells were grown in Advanced Dulbecco’s Modified Eagle Medium (ADMEM, Gibco) supplemented with 10 % FBS (fetal bovine serum, Biowest) and L-glutamine (Gibco) in 5% CO2 at 37°C. Cells were kept in culture up to passage number 15.

### Germ-free mice experiments

C57/BL6 6-week-old gnotobiotic male and female mice from Institut Pasteur Animal facilities (Janvier Labs) were acclimated on independent isolators (one isolator per strain) for a week prior to *C. difficile* challenge. Later mice were challenged with *C. difficile* spores (2x10^3^ per mice) by oral gavage. Mice health was monitored daily as described previously^98^. Progression of disease was assessed via Body Condition Scoring and body mass measurements^99^. Mice were followed to 13 days post *C. difficile* challenge.

### Transwell intestinal model (TIM)

Caco-2 cells or Caco-2-HT29-MTX co-culture, were seeded into 12-well Transwell inserts (pore size 0.4 µm, Corning) at a density of 2x10^5^ cells/cm^2^ and cultured for 18 days at 5% CO2 at 37°C. Cell culture media was changed three times a week.

### Intestine-on-a-chip model (IoC)

IoC-associated instrumentation and software were obtained from Emulate (Human Emulation System, Boston MA). Chips were prepared following manufacturer instructions and as described previously^100,101^. Briefly, chips were activated using ER1/ER2 solution (Emulate, 0.5 mg/mL) under UV for 20 minutes (36W, 365 nm) then washed once with ER2 solution (Emulate), followed by 2 PBS washes (Gibco). Chips were coated overnight at 4°C with ECM composed of 200 ug/mL of human Collagen IV (Sigma) + 100 ug/mL of Matrigel (Corning). ECM materials were washed twice with PBS followed by cell culture media. Caco2/TC7 cells were added to the epithelial channel at a concentration of 10^6^ cells/mL density. Caco2/TC7 and HT29-MTX co-culture was prepared by adding 8x10^5^ cells/mL of Caco2 + 2x10^5^ cells/mL of HT29- MTX (4:1 ratio). Cells were incubated for 1 day under static conditions at 37 °C with 5% CO2. After the cells adhere to the substrate, chips were gently washed with warm cell culture media to remove non-attached cells, then connected to the primed Pods (Emulate). Pods-chips were kept in the Zoë (Emulate) with a flow of 30 uL/h on the top and bottom channels for 1 day, then adding a stretch (10 %, 0.15 Hz) for 6 days. Cell culture medium reservoirs were refilled every 3 days.

### Lactate dehydrogenase release assays

To measure Lactate dehydrogenase release from Caco-2 or Caco-2 co-cultured with HT29-MTX in the TIM or IoC models, we used the commercial kit CytoTox 96 Non-Radioactive Cytotoxicity Assay (Promega) according to manufacturer instructions. The relative cytotoxicity obtained from cultures under normoxia conditions (at 5% CO2) was considered as 0% and these values were compared to hypoxia conditions over time (4% O2 and 5% CO2).

### TIM infection under hypoxia

*C. difficile* strains were cultured overnight (ON) on TY broth, the next day ON cultures were diluted (1:50) with new fresh media to obtain exponential phase bacteria (*λ*600nm 0.3 to 0.5). Bacteria were diluted to 10^6^ bacteria/mL in equilibrated ADMEM before infection. Cells were equilibrated 1h before infection under hypoxia conditions (4% O2, 5% CO2), then wells were infected with 500 uL of bacterial suspension (10^6^ bacteria/mL).

### TIM adhesion assays

Infection conditions were kept as indicated previously for TIM infection under hypoxia conditions (4% O2, 5% CO2). After 3, 6, 18 or 24 h of incubation cells were washed three times with 500uL of PBS (Gibco) to eliminate non-adherent bacteria. Cells and adherent bacteria were diluted in 500 uL of PBS (Gibco) and recovered by scraping the Transwell wells with 1mL tips and centrifugation (5 min at 5 000 rpm). Adherent bacteria were serially diluted and plated on TY agar plates, incubated for 48 h at 37°C under anaerobic conditions.

### IoC infection under hypoxia

*C. difficile* strains were cultured overnight (ON) on TY broth, the next day ON cultures were diluted (1:50) with new fresh media to obtain exponential phase bacteria (*λ*600nm 0.3 to 0.5). Bacteria were diluted to 10^6^ bacteria/mL in equilibrated ADMEM before infection. IoC were equilibrated 6h before infection by decreasing O2 levels each hour (18%, 15%, 12%, 9%, 6%, 4%; with 5% CO2) in the housing cell culture incubator. IoC chips were disconnected from the pods and infected under static conditions (no flow, no stretch) with 50 uL of bacterial suspension (10^6^ bacteria/mL). After 1h 30 min, chips were reconnected to the Pods and reintroduced in the Zoë with a flow of 30 μL/h (no stretch during infection).

### TIM and IoC immunostaining

TIM and IoC were fixed with 4% of paraformaldehyde (Electron Microscopy Sciences) diluted in PBS with Ca^2+^ and Mg^2+^ (Gibco) for 30 min. After fixation Transwell and chips were washed three times with PBS and stored at 4°C. For the IoC transversal sections, chips were cut in 300- µm thick slices using a vibrating blade microtome (VT1000S, Leica). IoC sections and Transwell were permeabilized with 0.1% Triton X-100 in PBS with Ca^2+^ and Mg^2+^ (Gibco) for 20 min at room temperature (RT) and then washed three times with PBS. Later, blocking solution (2% BSA in PBS with Ca^2+^ and Mg^2+^) was added for 1h at RT.

### Spinning disk fluorescence microscopy

Images were performed in a Nikon Ti-E inverted microscope equipped with a Perfect Focus System (TI-ND6-PFS Perfect Focus Unit) and a Yokogawa confocal spinning disk unit (CSU-W1) using a 60X/1.42 NA oil objective. A Z-stack of 300 to 800 planes with 0.3 µm z-steps was acquired sequentially in 4 channels (Da/Fi/Tr/Cy5-4x-B, Finkel Quad FF01-440/521/607/700).

### Production of LMW-SlpA specific monoclonal antibodies NF10 and QD8

Knock-in mice expressing human antibody variable genes for the heavy (VH) and kappa light chain (Vκ) were previously described ^102,103^ and provided by Regeneron Pharmaceuticals to be bred at Institut Pasteur. BALB/c mice were purchased from Janvier Labs. All animal care and experimental procedures were conducted in compliance with national guidelines. The study, registered under #210111, was approved by the Animal Ethics Committee of CETEA (Institut Pasteur, Paris, France) and by the French Ministry of Research. BALB/c and VelocImmune mice were injected intraperitoneally on days 0, 21, and 42; with 50 μg of either recombinant LMW630 mixed with 200 ng/mouse pertussis toxin (Sigma-Aldrich, MO, USA) for NF10 production or with 50 μg of each of five recombinant LMWs in alum mixed with 200 ng/mouse pertussis toxin (Sigma-Aldrich, MO, USA) for QD8 production. An enzyme-linked immunosorbent assay (ELISA) previously described^35^, was performed to measure serum responses to antigens and the three immunized animals with the highest serum titers were boosted with the same preparation. Four days later, splenocytes were fused with myeloma cells P3X63Ag8 (ATCC, France) using a ClonaCell-HY Hybridoma Kit, according to the manufacturer instructions (StemCell

Technologies, Canada). Culture supernatants were screened using ELISA^35^, and antigen-reactive clones were expanded in serum IgG-free RPMI-1640 (Sigma-Aldrich) into roller bottles at 37°C. After 14 days, the supernatants were harvested by centrifugation at 2 500 rpm for 30 min and filtered through a 0.2 μm filter. Antibodies were purified by Protein A affinity chromatography (AKTA, Cytiva, Germany), as described previously ^104^.

### Minimal Inhibitory Concentration (MIC) determination and antibiotic resistance assays

MICs were determined by broth microdilution as described before^105^. Briefly, a 96-well plate containing twofold dilutions of desired antibiotic were inoculated with ON culture diluted to a final *λ*600nm of 0.05 in ADMEM (Gibco) supplemented with 10 % FBS (Biowest) and L-glutamine (Gibco). After 24 h at 37 °C, MIC was determined by measuring λ600nm in a plate reader (Promega GloMax Explorer). Supplemented ADMEM medium was used as a blank.

For the antibiotic resistance assays in TIM, infections were performed as indicated previously and 24h p.i, antibiotics were added at 1x, 10x and 100x the MIC. MIC for fidaxomicin was defined as 1 μg/mL and for vancomycin as 12.5 μg/mL. After 24h of antibiotics treatment, resistant bacteria were recovered by scraping the Transwell wells with 1mL tips and centrifugation (5 min at 5 000 rpm). Resistant bacteria were serially diluted and plated on TY agar plates, incubated for 48 h at 37°C under anaerobic conditions.

### *In vitro* biofilm assays

*C. difficile* ON cultures were diluted to a final *λ*600nm of 0.02 into fresh equilibrated Gut Microbiota Medium (GMM)^46^ or GMM supplemented with mucin, 1 mL per well was deposited in 24-well polystyrene tissue culture-treated plates (Falcon Clear Flat Bottom) and the plates were incubated at 37 °C in anaerobic environment for 48h. Type II mucin (Sigma M2378) and native mucin extracted from pork were diluted in Milli-Q water (concentration 40mg/mL), autoclaved (15 min, 121°C) and added to the pre-equilibrated medium (final concentration 2 mg/mL). Biofilm biomass was measured using established methods^42^. Briefly, spent media was removed. Biofilms were air dried and stained with crystal violet (CV; 0.2% w/v) for 10 min. CV was removed by inversion; wells were washed twice with PBS and then air-dried. Dye bound to the biofilm biomass was solubilized by adding 1 mL of 75% (%v/v) ethanol and the absorbance, corresponding to the biofilm biomass, was measured at a *λ*600nm with a plate reader (Promega GloMax Explorer). When needed, the solubilized dye was diluted with 75% ethanol for the reading to remain in the linear range. Sterile GMM or GMM with mucin was used as a blank for the assays.

### CDT toxin assays

The two CDT subunits, CDTa and activated CDTb, were generated, as previously described, using an *E. coli* expression system ^39^. Briefly, the complete ORFs of CDTa and CDTb were amplified by PCR from genomic DNA of *C. difficile* strain R20291 (GenBank: FN545816.1). Only the sequences bp 127–389 for CDTa and bp 127–2628 for CDTb (without the leader sequences) were cloned into the pGEX-2T vector to genetically engineer the GST fusion proteins of the mature CDTa and CDTb. GST– CDTa and GST–CDTb were expressed in *E. coli* following a standard protocol. Gene expression was induced by 100 μM isopropyl-β-D-thiogalactopyranosid when the bacterial cultures reached an OD600nm of 0.6. The GST fusion proteins were affinity purified via glutathione-sepharose (GE Healthcare, Dornstadt, Germany) by gravity flow, and the proteins were released either by thrombin (0.06U/μg protein, 4 ◦C overnight for CDTa) or by elution with 10 mM glutathione (CDTb). Eluted GST–CDTb was directly activated by trypsin (0.2 μg/μg protein, 30 min at RT). Trypsin was inactivated by 2 mM 4-(2-Aminoethyl) benzensulfonylfluorid, and the solution was dialyzed against PBS ON.

### ELISA-based measurement of CDT

A 96-well immuno-plate (Nunc Maxisorp) was coated ON with CdtB capture antibody (MBS396782, MyBioSource) diluted into PBS. Plates were washed twice (PBS + 1% Tween 20). Blocking buffer (PBS + 2% BSA) was added and plates were incubated for at least 1 hour at RT and washed twice. Bacterial supernatants or lysates were serially diluted in PBS and incubated in coated plates for 90 min at RT. After two washes, chicken anti-CdtB IgY HRP conjugated antibody (MBS396785, MyBioSource) was added for 1-2 h at RT. The wells were washed four times and incubated with TMB (3,3′,5,5′tetramethylbenzidine) HRP substrate solution (Thermo Fisher Scientific) for 5 to 30 min in the dark. The stop solution (H2SO4; 0.2 M) was added into each well and the absorbance of the reaction was read at 450 nm (Promega Glomax Explorer plate reader).

### ELISA-based measurement of TcdA

Total TcdA amount was quantified from supernatants. Briefly, 1.5 mL of culture was harvested by centrifugation for 4 min at 13 000 rpm. Supernatants were collected and bacterial pellets were frozen at −20 °C. The supernatants fractions were then analyzed by ELISA. A 96-well immuno-plate (Nunc Maxisorp) was coated with 2 μg/mL of anti- toxin A rabbit polyclonal antibody (Abcam, Inc.) ON at 4 °C. The coated wells were washed and incubated with Superblock blocking buffer (Thermo Fisher Scientific) for 1 h. The wells were then washed and air-dried. Samples were added into the wells, and the plate was incubated at 37 °C for 90 min. After washings, 0.2 μg/mL of an anti- toxin A chicken horseradish peroxidase (HRP) antibody (LSBio) was added in each well and the plate was incubated for 1 h at 37 °C. The wells were washed and incubated with a TMB (3,3′,5,5′tetramethylbenzidine) substrate solution (Thermo Fisher Scientific) for 15 min in the dark. The stop solution (H2SO4; 0.2 M) was added into each well and the absorbance of the reaction was read at 450 nm (Promega Glomax Explorer plate reader).

### RNA isolation and quantitative reverse-transcriptase PCR

Cells were washed once with PBS (Gibco), lysed in RLT buffer (Qiagen) and freeze at -80°C until extraction was performed. RNA was extracted with RNeasy mini Kit (Qiagen) following manufacturer recommendations. DNA digestion was carried out in columns using RNase-free DNase set (Qiagen) and RNA clean-up with a RNeasy MinElute Cleanup kit (Qiagen). The RNA yield was measured with Nanodrop. cDNA was obtained with QuantiTect Reverse Transcription Kit (Qiagen) following manufacturer instructions. The quantitative Real-Time PCR was performed on StepOne Real-Time PCR Systems (Thermo Scientific) using SsoFast EvaGreen Supermix (Bio-Rad) following manufacturer instructions. Each reaction was performed in technical triplicate with 2 or 3 independent biological replicates. Data were analyzed by the ΔΔCt method. Gene expression levels were normalized to the *rps13* gene.

### Spore preparation

Spore suspensions were prepared as previously described^106^. Briefly, 200 μl from ON cultures of *C. difficile* strains were plated on sporulation medium for *Clostridioides difficile* (SMC) medium (9% Bacto peptone, 0.5% proteose peptone, 0.15% tris base, and 0.1% ammonium sulfate) and were incubated at 37°C for 7 days under anaerobic conditions. Spores were scraped off and resuspended in 2 mL of sterile ice cold water and incubated for 7 days at 4°C. Cell fragments and spores were separated by centrifugation using a HistoDenz (Sigma-Aldrich) gradient^107^. Spores were enumerated on TY supplemented with 1% taurocholate and kept at 4°C on glass vials.

### Ethics statement

Animal studies were performed in agreement with European and French guidelines (Directive 86/609/CEE and Decree 87-848 of 19 October 1987). The study received the approval of the Institut Pasteur Safety Committee (Protocol n°18086) and the ethical approval of the local ethical committee “Comité d’Ethique en Experimentation Animale Institut Pasteur no. 89 (CETEA)” (CETEA dap190131).

### Germ-free mice infection experiments

C57/BL6 7-week-old gnotobiotic male and female mice from Institut Pasteur Animal facilities (Janvier Labs) were challenged with *C. difficile* spores (2x10^3^ per mice) by oral gavage. To assess bacterial persistence, fecal pellets were collected over a 13- day period (days 0, 1, 2, 6, 7, 8, 9 and 13). Fecal pellets were homogenized in the anaerobic hood in 1mL of PBS, serially diluted, and plated in triplicate on BHI agar containing 2% defibrinated horse blood, 0.1% taurocholate, tetracycline (5 μg/mL), ciprofloxacin (5 μg/mL) cefoxitin (8 μg/mL), and cycloserine (250 μg/mL) to assess the total number of CFUs. To assess the total number of spores, diluted fecal pellets were incubated in ethanol (50% v/v final concentration) for at least 1h and plated in triplicates using the same medium.

### Measurement of lipocalin-2 intestinal levels

Frozen fecal samples were reconstituted in PBS and vortexed for 5 min to homogenize the fecal suspension. Then samples were centrifuged for 10 min at 10 000 rpm and 4°C. Clear supernatants were collected and stored at −20°C until analysis. Lcn-2 levels were estimated in the supernatants using Duoset murine Lcn-2 ELISA kit (R&D Systems). Samples from day 0 (before infection) were used as negative controls.

### Histological processing and staining of tissue samples

Intestinal tissues were recovered, and full rolls were placed in Carnoy’s fixative solution (60% ethanol, 30% chloroform, and 10% glacial acetic acid) ON at 4°C. Later, ethanol gradients were applied to wash fixed tissues (70%,80%, 95% and 100 % vol/vol). Tissues were embedded in ethanol/xylene (1:1) and xylene, followed by embedding in Paraffin. Tissue blocks were laterally sectioned at 10 μm and were stained with hematoxylin and eosin (H&E) to asses histological score or perform immunostainings.

### Measurement of mucin thickness and goblet cells in the colon of infected mice

Colonic sections were also stained with Alcian Blue, preferentially staining mucopolysaccharides, and 40 crypts were randomly selected per animal to determine goblet cell number per crypt.

### Image analysis of *C. difficile* biofilms in the TIM and IoC models

TIM and IoC images were analyzed using the same analysis scripts, developed in Python^108^. First, 3D images were projected along Z to yield 2D multi-channel images. For each image, chromatic aberration was corrected by registering the bacteria channel with respect to the mucin channel, using phase cross-correlation^109^ implemented in scikit-image^110^. The mucin signal was quantified by first segmenting the tissue surface in the image, on the nuclei channel combined with the actin channel, using an intensity threshold. The mean mucin signal and its standard deviation were then measured within the resulting tissue mask. Bacteria were segmented in the far-red channel using Omnipose^111^. A bacterium was classified as positive for mucin if the mean mucin intensity within the bacteria mask was larger than the mean mucin signal in the tissue plus the standard deviation. The count of all mucin-positive and negative bacteria and their total surface were then reported for each image. Results were exported to ImageJ TIFFs and ImageJ ROIs with the tifffile tool^112^ and were manually inspected using Fiji^113^.

### Image analysis of *C. difficile* biofilms in the colon and cecum of infected mice

Colon and caecum images were analyzed like TIM and IoC models with minor modifications (see above). Bacteria were segmented as a mask and not as single bacteria by thresholding after filtering by a 9x9 median filter and a gaussian filter with σ=0.5 pixels.

### Statistical analysis

Statistical significance was determined using unpaired *t* tests or multiple unpaired Holm-Sidak t tests. For multiple comparisons, analysis of variance (ANOVA) was used with Bonferroni’s, Dunnett’s or Geisser-Greenhouse correction as recommended. Mann Whitney tests were performed for biofilms, mucin thickness and goblet cells analyses. Statistics were completed using Prism 8.0 (GraphPad Software). Specific details with regard to statistical tests, statistical significance values (“p”), sample sizes (‘‘n’’) and replicates are indicated in the figure legends. For all analysis, significance was considered as p<0.05.

### Data and code availability

The full code for the image analysis performed in this paper is available publicly at https://gitlab.pasteur.fr/iah-public/clostridioides-difficile-binary-toxin-cdt-induces-biofilm-like-persisting-microcolonies

## ACKNOWLEDGMENTS

We thank Nathalie Sauvonnet and Valérie Malarde for their great support and advices with transwell and organ on chip models. We also thank Ralf Gerhard for providing us the purified CDT toxin. We thank Martine Jacob, Céline Mulet, and Thierry Pedron for their technical support with the mice experiments. We thank David Hardy, Sabine Maurin, Magali Tichit and Mélanie Juchet-Martin for their expertise and technical support with histological experiments. We thank Jost Enninga and Laurent Audry for their help and advices with spinning disk microscopy. Funding: This work was supported by the Institut Pasteur, the “Integrative Biology of Emerging Infectious Diseases” (LabEX IBEID) funded in the framework of the French Government’s “Programme Investissements d’Avenir” and the Agence National de la Recherche in the framework of the: ANR-20-CE15-0022 (DifBioRel) to J.M.T. and B.D., and ANR- 20-CE15-0003 (Difficross) to J.P. Institut Carnot Pasteur Microbe & Santé supported Emulate chips purchase. We acknowledge the France-BioImaging infrastructure supported by the French National Research Agency (ANR-10-INBS-04).

## AUTHOR CONTRIBUTIONS

Conceptualization J.M.T and B. D.; Investigation J.M.T. (Transwell and organ on chip model’s standardization, mice and infection experiments, immunostaining, microscopy, and qRTPCR); B.C. (histological analysis); H.M, M.K. (Organ on chip seeding and maintenance); J.P., P.A.S., E.L. (mutants); S.C.R., A.C. (*in vitro* biofilm, ELISAS and growth curves); L.H. (production of monoclonal antibodies); Data analyses J.M.T., B.D., B.C.; Image analysis J.Y.T.; Writing-Original Draft J.M.T. and B.D.; Writing–Review & Editing J.M.T., B.D., B.C., J.P., S.G., J.Y.T.; Supervision J.M.T and B. D.

