## Supplemental Data for "*Clostridioides difficile* binary toxin CDT induces biofilm-like persisting microcolonies"

### SUPPLEMENTAL INFORMATION

**Table S1. Strains and plasmids used in this study.**

| Strain | Genotype | Origin |
|---|---|---|
| ***E. coli*** | | |
| NEB-10 beta | *Δ(ara-leu) 7697 araD139 fhuA ΔlacX74 galK16 galE15 e14- ϕ80dlacZΔM15 recA1 relA1 endA1 nupG rpsL* ($Str^R$) *rph spoT1 Δ(mrr-hsdRMS-mcrBC)* | New England Biolabs |
| HB101(RP4) | *supE44 aa14 galK2 lacY1* Δ(*gpt-proA*) 62 *rpsL20* ($Str^R$)*xyl-5 mtl-1 recA13* Δ(*mcrC-mrr*) *hsdS*$_B$ ($r_B$-$m_B$-) RP4 ($Tra^+$ IncP $Ap^R$ $Km^R$ $Tc^R$) | Laboratory stock |
| ***C. difficile*** | | |
| 630Δ*erm* | 630Δ*erm* 012 | Reference[1] |
| CD1611 | *630Δerm* $ToxAB^-$ | This study |
| CD1666 | UK1 | Reference[2] |
| CNRS_CD129 | UK1 $ToxAB^-$ $CDT^+$ | This study |
| CNRS_CD345 | UK1 $ToxAB^-$ $CDT^-$ | This study |
| **Plasmid** | | |
| pMSR0 | Allele exchange for C. *difficile* UK1 | Reference[3] |
| p127 | pMSR derivative for CdtLoc deletion | This study |
| p128 | pMSR derivative for PaLoc deletion | This study |

**Table S2. Oligonucleotides used in this study.**

| Primer | Sequence (5' to 3')* | Use |
|---|---|---|
| PA_001 | ttttttgttaccctaagtttCATGATATGGAAATTGCTG | 5' left arm for *cdtAB* deletion |
| PA_002 | gagtaattgcCATTTATTCTCCCTCCCAATATTAG | 3' left arm for *cdtAB* deletion |
| PA_003 | agaataaatgGCAATTACTCCAGACGATAG | 5' right arm for *cdtAB* deletion |
| PA_004 | agattatcaaaaaggagtttGCAGAAAAAGCCGAAAAAC | 3' right arm for *cdtAB* deletion |
| PA_005 | GTGATGGATTATGGATAGC | 5' *cdtAB* deletion screening |
| PA_006 | TGTGGGGACAAATTTAAATC | 3' *cdtAB* deletion screening |
| PA_007 | ttttttgttaccctaagtttGTTTGTTTTAGCAAGAAATAACTCAG | 5' left arm for *tcdBEA* deletion |
| PA_008 | tattttagccCATAAAATTTTCTCCTTTACTATAATATTTTTATTG | 3' left arm for *tcdBEA* deletion |
| PA_009 | aaattttatgGGCTAAAATATATGTTTGACAAAAAATTATTC | 5' right arm for *tcdBEA* deletion |

| | | |
|---|---|---|
| PA_010 | agattatcaaaaaggagtttCTTGTTCTGAAGACC ATG | 3’ right arm for *tcdBEA* deletion |
| PA_011 | GAGAGGATGATTTTATGC | 5’ *tcdBEA* deletion screening |
| PA_012 | CCATACCAGGGATAGCTGTAG | 3’ *tcdBEA* deletion screening |
| QRTBD014 | ATAAATTGCATGTTGCTTCATAACT | Intact PaLoc in wild type strains |
| OS234 | AGCTTTCGCTTTAGGCAGTG | Intact PaLoc in wild type strains |
| Muc5AC Fw | TGATCATCCAGCAGCAGGGCT | qPCR |
| Muc5AC Rv | CCGAGCTCAGAGGACATATGGG | qPCR |
| Muc1c Fw | ACTACTACCAAGAGCTG | qPCR |
| Muc1c Rv | CTCATAGGATGGTAGGT | qPCR |
| Muc2 Fw | ACTGCACATTCTTCAGCTGC | qPCR |
| Muc2 Rv | ATTCATGAGGACGGTCTTGG | qPCR |
| Rps13 Fw | CGAAAGCATCTTGAGAGGAACA | qPCR |
| Rps13 Rv | TCGAGCCAAACGGTGAATC | qPCR |

*Lowercase bases indicate overlapping sequences

**Table S3. Susceptibility of *C. difficile* strains to vancomycin and fidaxomicin cultured in supplemented ADMEM medium**

| Strains | Antibiotics | Reported MIC for WT (µg/mL) | MIC ADMEM (µg/mL) | Reference |
|---|---|---|---|---|
| UK1 WT<br>UK1 ToxAB$^{-}$ CDT$^{+}$<br>UK1 ToxAB$^{-}$ CDT$^{-}$ | Vancomycin | 1 | 12.5 | [4] |
| UK1 WT<br>UK1 ToxAB$^{-}$ CDT$^{+}$<br>UK1 ToxAB$^{-}$ CDT$^{-}$ | Fidaxomicin | 0.25 | 1 | [5] |

### SUPPLEMENTAL FIGURES

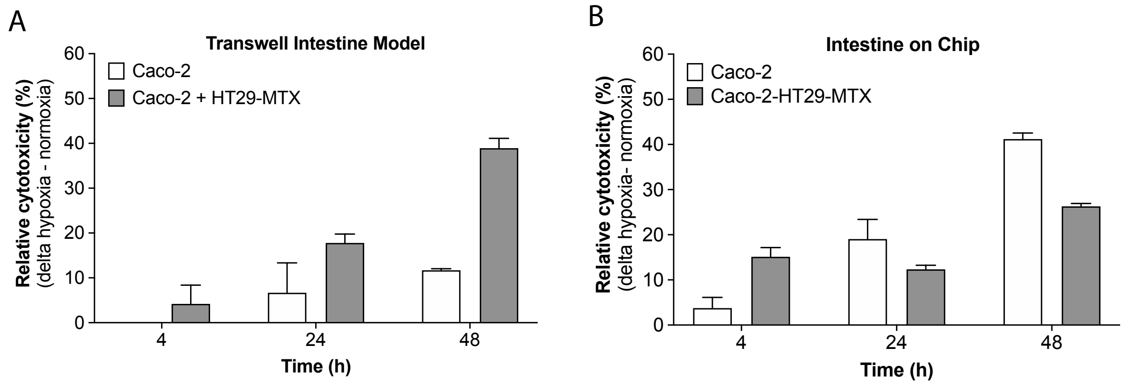


**Figure S1. Cell survival under hypoxia in the Transwell Intestine model and Intestine-on chip model.** Lactate dehydrogenase release assays were performed to evaluate cell survival in (A) the Transwell intestine model (TIM) and (B) the Intestine-on-chip (IoC) model after 4, 24 and 48h under normoxia or hypoxia (4% $O_2$, 5% $CO_2$). The relative cytotoxicity is represented as the difference between the hypoxia and normoxia conditions (delta hypoxia-normoxia). Data represents mean with SEM. Data represents mean with SEM (C, D, E).

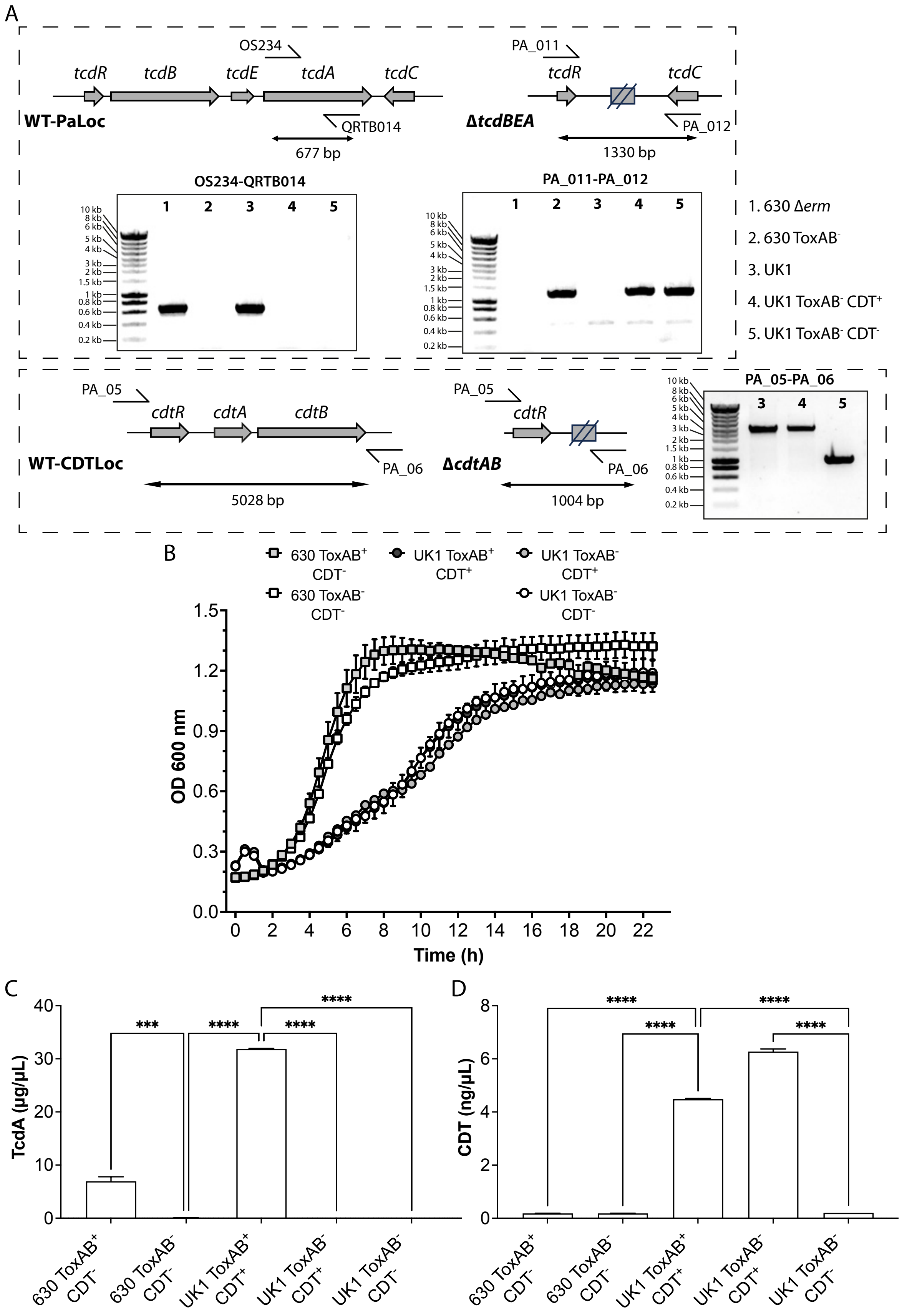
A
OS234
tcdR
tcdB
tcdE
tcdA
tcdC
WT-PaLoc
QRTB014
677 bp
PA_011
ΔtcdBEA
PA_012
1330 bp
OS234-QRTB014
PA_011-PA_012
1 2 3 4 5
10 kb
8 kb
6 kb
5 kb
4 kb
3 kb
2 kb
1.5 kb
1 kb
0.8 kb
0.6 kb
0.4 kb
0.2 kb
1. 630 Δerm
2. 630 ToxAB-
3. UK1
4. UK1 ToxAB- CDT+
5. UK1 ToxAB- CDT-
PA_05
cdtR
cdtA
cdtB
WT-CDTLoc
PA_06
5028 bp
ΔcdtAB
1004 bp
PA_05-PA_06
3 4 5
B
630 ToxAB+ CDT-
UK1 ToxAB+ CDT+
UK1 ToxAB- CDT+
630 ToxAB- CDT-
UK1 ToxAB- CDT-
OD 600 nm
Time (h)
C
TcdA (μg/μL)
D
CDT (ng/μL)
630 ToxAB+ CDT-
630 ToxAB- CDT-
UK1 ToxAB+ CDT+
UK1 ToxAB- CDT+
UK1 ToxAB- CDT-

**Figure S2. Growth of generated PaLoc and CdtLoc *C. difficile* mutants and absence of TcdA and CDT toxins in culture supernatants A.** (A) PCR verification of the *tcdBEA* deletion mutants in both 630 Δ*erm* and UK1 strains and PCR verification of the *cdtAB* deletion mutants in derivative *tcdBEA* mutant of *C. difficile* UK1 strain. The pair OS234-QRTB014 that amplify an internal fragment of *tcdA* gene was used to verify the intact PaLoc, PA_011-PA_012 to verify the *tcdBEA* deletion and PA_05-PA_06 to verify the *cdtAB* deletion. (B) Growth curves of 630 Δ*erm*, UK1 WT or *tcdBEA* and *cdtAB* mutant strains in TY medium supplemented with glucose. (C) TcdA toxin or (D) CDT toxin secretion in extracellular fractions of 630 Δ*erm*, UK1 WT or *tcdBEA* and *cdtAB* mutant strains after 24h of growth in TY medium. A one-way ANOVA was performed and statistical significance is represented (***p <0.001 and **** p<0.0001). Data represents mean with SEM (C, D, E).

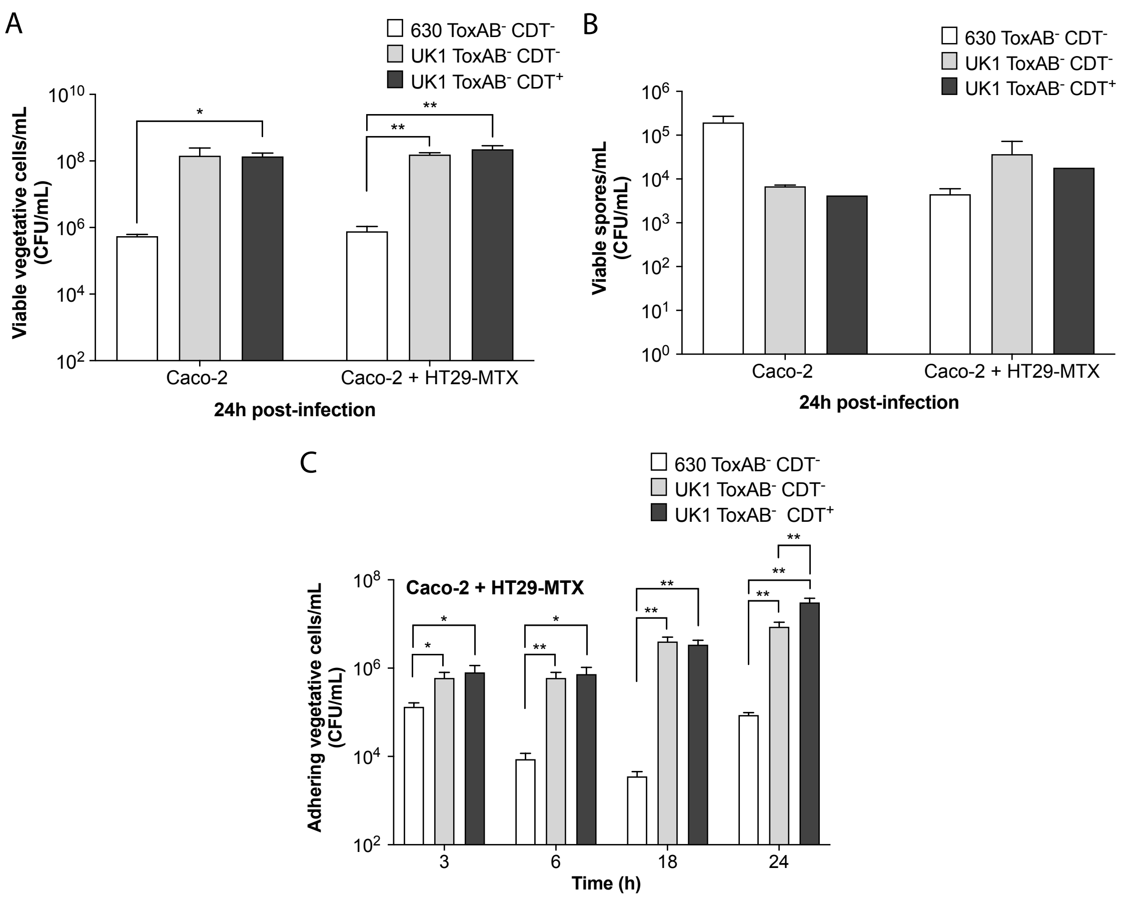


**Figure S3. Growth and adhesion of *C. difficile* strains in the Transwell Intestine model 24h after infection.** Caco-2 cells alone or co-cultured with HT29-MTX cells in the TIM model were infected with 630 ToxAB$^{-}$CDT$^{-}$, UK1 ToxAB$^{-}$CDT$^{-}$ and UK1 ToxAB$^{-}$CDT$^{+}$ strains. (A) Total viable vegetative cells (CFU) were numbered 24h p.i. (B) Adhering vegetative cells were numbered 24h p.i after eliminating non-adhering vegetative cells by PBS washes. Data represents mean with SD (A, C) or SEM (B). Multiple unpaired *t* tests were performed and statistical significance is represented (* p<0.05, ** p <0.01).

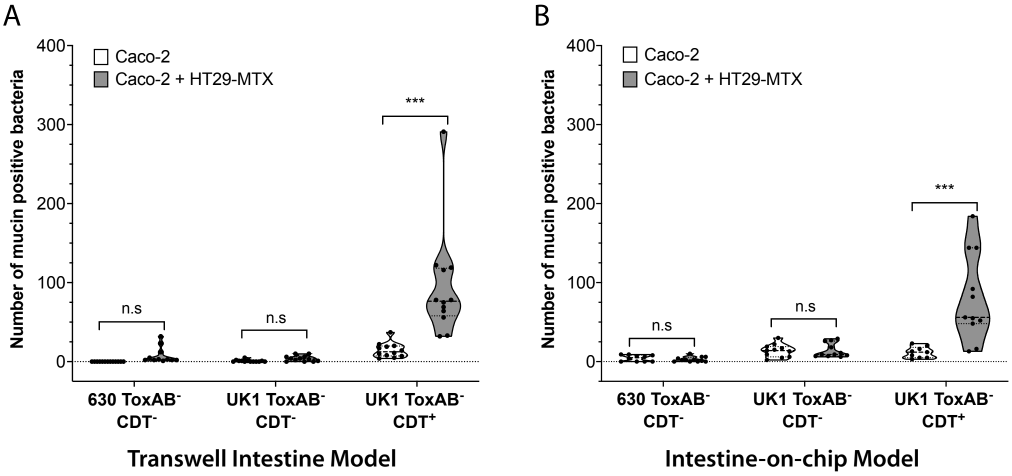


**Figure S4. CDT-dependent microcolonies co-localize with mucin in a Transwell Intestine model and Intestine-on-chip models.** Caco-2 alone or cocultured with HT29-MTX cells were infected with 630 $ToxAB^{-}CDT^{-}$, UK1 $ToxAB^{-}CDT^{-}$, or UK1 $ToxAB^{-}CDT^{+}$ under hypoxic conditions (4% $O_2$, 5% $CO_2$). Total mucin positive bacteria counted in Caco-2 cells alone or with HT29-MTX cells infected in (A) the TIM at 24h p.i or (B) the IoC model at 48h p.i with *C. difficile* strains as indicated. The number of mucin positive bacteria are reported for each image and at least 10 images were quantified. Each black square in the graph represents one image. Data and quantifications are representative of 2 (IoC) or 3 (TIM) independent biological replicates. Multiple unpaired *t* tests were performed and statistical significance is represented with ***p <0.001.

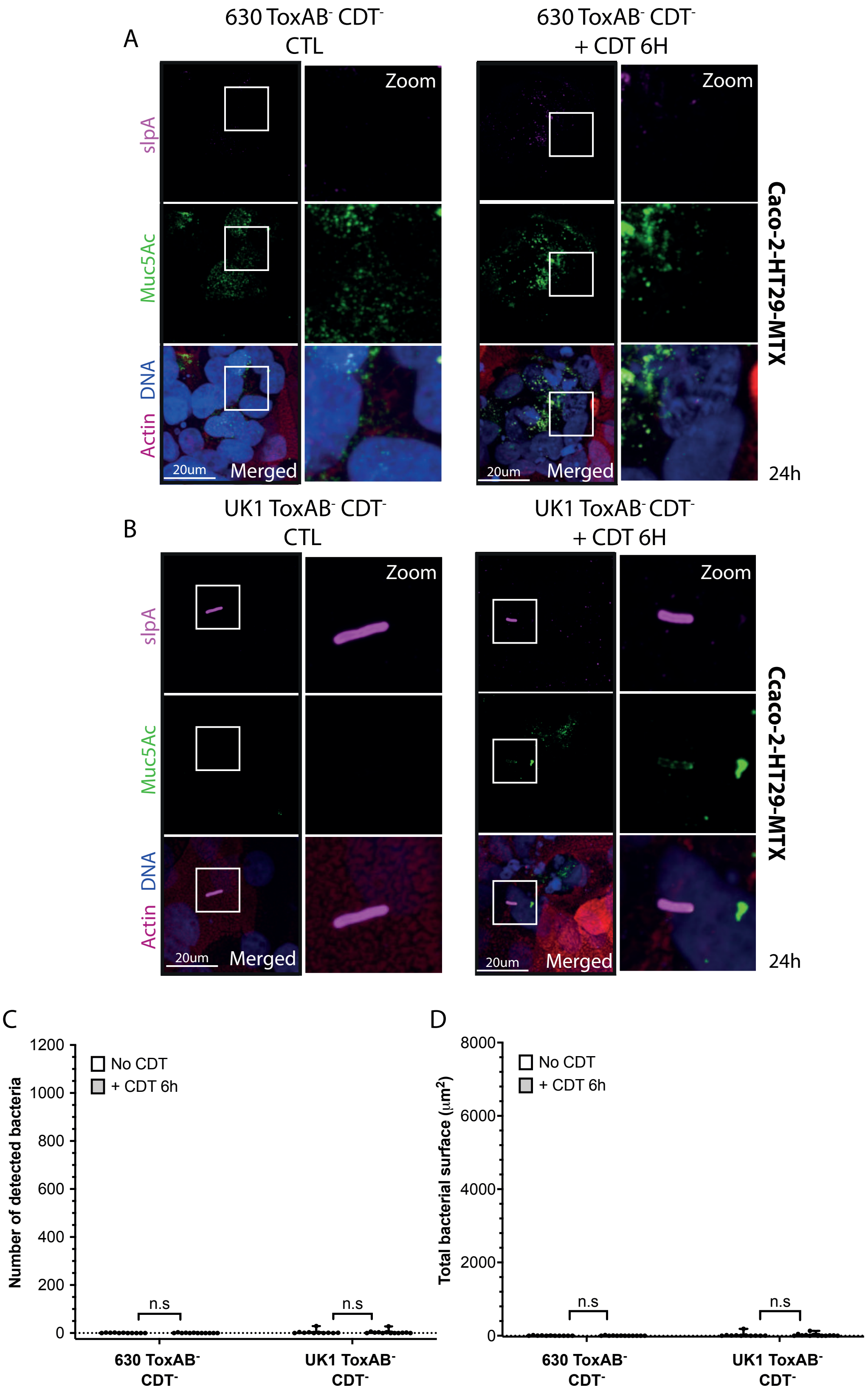
A
630 ToxAB- CDT-
CTL
630 ToxAB- CDT-
+ CDT 6H
Zoom
slpA
Muc5Ac
Actin DNA
20um
Merged
Caco-2-HT29-MTX
24h
B
UK1 ToxAB- CDT-
CTL
UK1 ToxAB- CDT-
+ CDT 6H
Ccaco-2-HT29-MTX
C
Number of detected bacteria
No CDT
+ CDT 6h
n.s
630 ToxAB- CDT-
UK1 ToxAB- CDT-
D
Total bacterial surface (µm2)

**Figure S5. $CDT^{-}$ strains treated with purified CDT toxin during 6h in in the Transwell intestine model.** Representative 3D reconstructed images of Caco-2 cocultured with HT29-MTX cells infected with (A) 630 $ToxAB^{-}CDT^{-}$ or (B) UK1 $ToxAB^{-}CDT^{-}$ during 24h under hypoxic conditions (4% $O_2$, 5% $CO_2$). Infected intestinal cells were exposed to CdtA (200ng/mL) and activated CdtB (400 ng/mL) during 6h. DNA was labelled with DAPI (blue), mucin with anti-Muc5AC AF488 (green), actin with phalloidin rhodamine (red) and *C. difficile* with anti-SlpA[6] AF647 (magenta). (C) Number of bacteria detected 24h p.i in Caco-2 cells cocultured with HT29-MTX cells infected with *C. difficile* strains as indicated. (D) Total bacteria surface detected 24h p.i in cells infected with *C. difficile* strains as indicated. The number of bacteria and total bacterial surface detected are reported for each image and at least 10 images were quantified per condition. Each black circle in the graph represents one image.

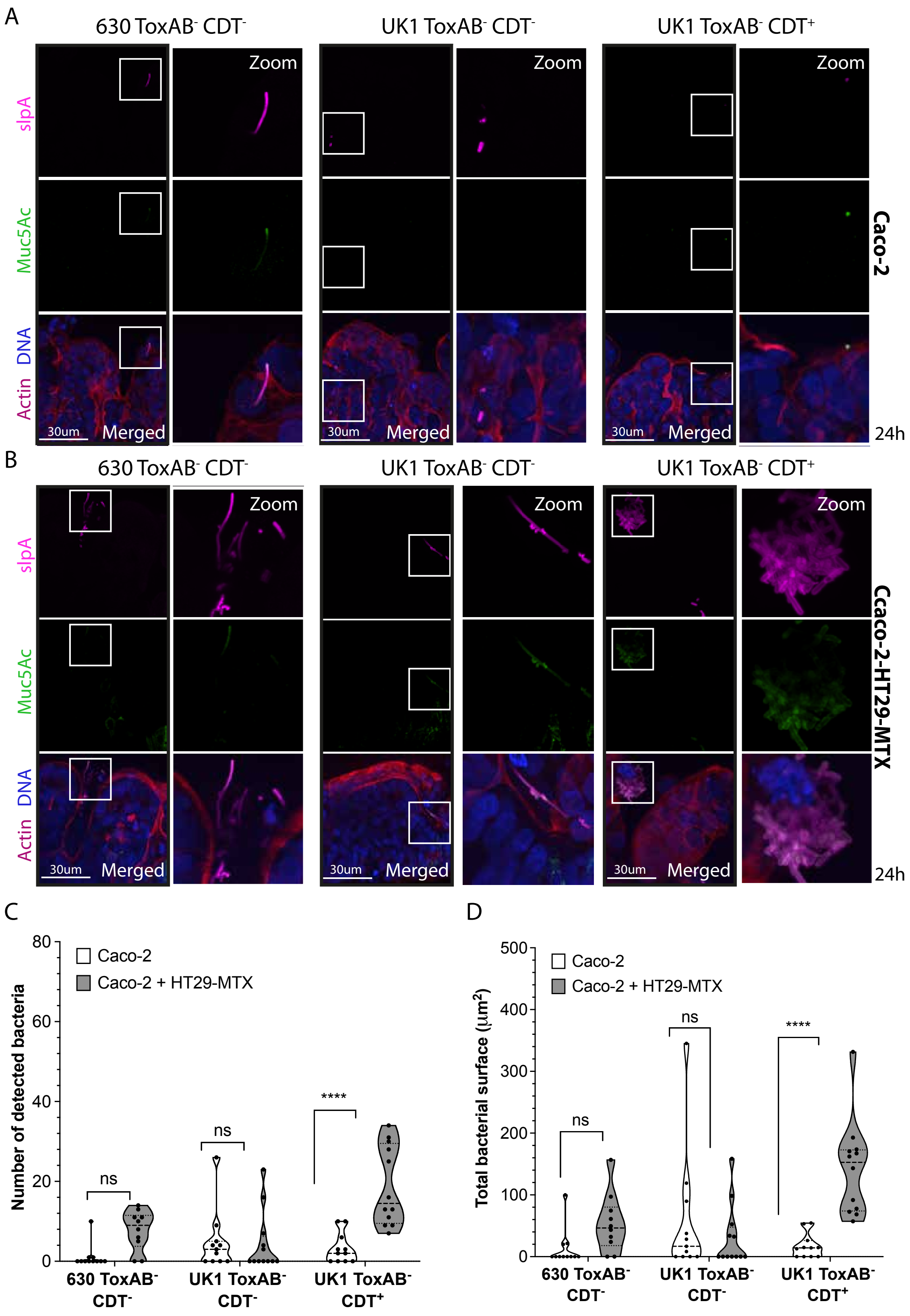
A
630 ToxAB- CDT-
UK1 ToxAB- CDT-
UK1 ToxAB- CDT+
Zoom
slpA
Muc5Ac
DNA
Actin
30um
Merged
Caco-2
24h
B
Ccaco-2-HT29-MTX
C
Caco-2
Caco-2 + HT29-MTX
Number of detected bacteria
ns
****
630 ToxAB- CDT-
UK1 ToxAB- CDT-
UK1 ToxAB- CDT+
D
Total bacterial surface (µm2)

**Figure S6. CDT$^{+}$ strain forms clumps in an Intestine-on-chip model 24h post-infection.** Representative 3D reconstructed images of (A) Caco-2 cells or (B) Caco-2 cells cocultured with HT29-MTX cells infected with 630 ToxAB$^{-}$CDT$^{-}$, UK1 ToxAB$^{-}$CDT$^{-}$ or UK1 ToxAB$^{-}$CDT$^{+}$ during 24h under hypoxic conditions as previously indicated. DNA was labelled with DAPI (blue), anti-Muc5AC AF488 (green), actin with phalloidin rhodamine (red) and *C. difficile* with anti-SlpA AF647 (magenta). (C) Number of bacteria detected 24h p.i in Caco-2 cells alone or with HT29-MTX cells infected with *C. difficile* strains as indicated. (C) Total bacteria surface detected 24h p.i in Caco-2 cells alone or with HT29-MTX cells infected with *C. difficile* strains as indicated. The number of bacteria and total bacterial surface detected are reported for each image and at least 10 images were quantified per condition. Each black circle in the graph represents one image. Data and quantifications are representative of 1 independent biological replicate. Multiple unpaired *t* tests were performed and statistical significance is represented with **** $p<0.0001$.

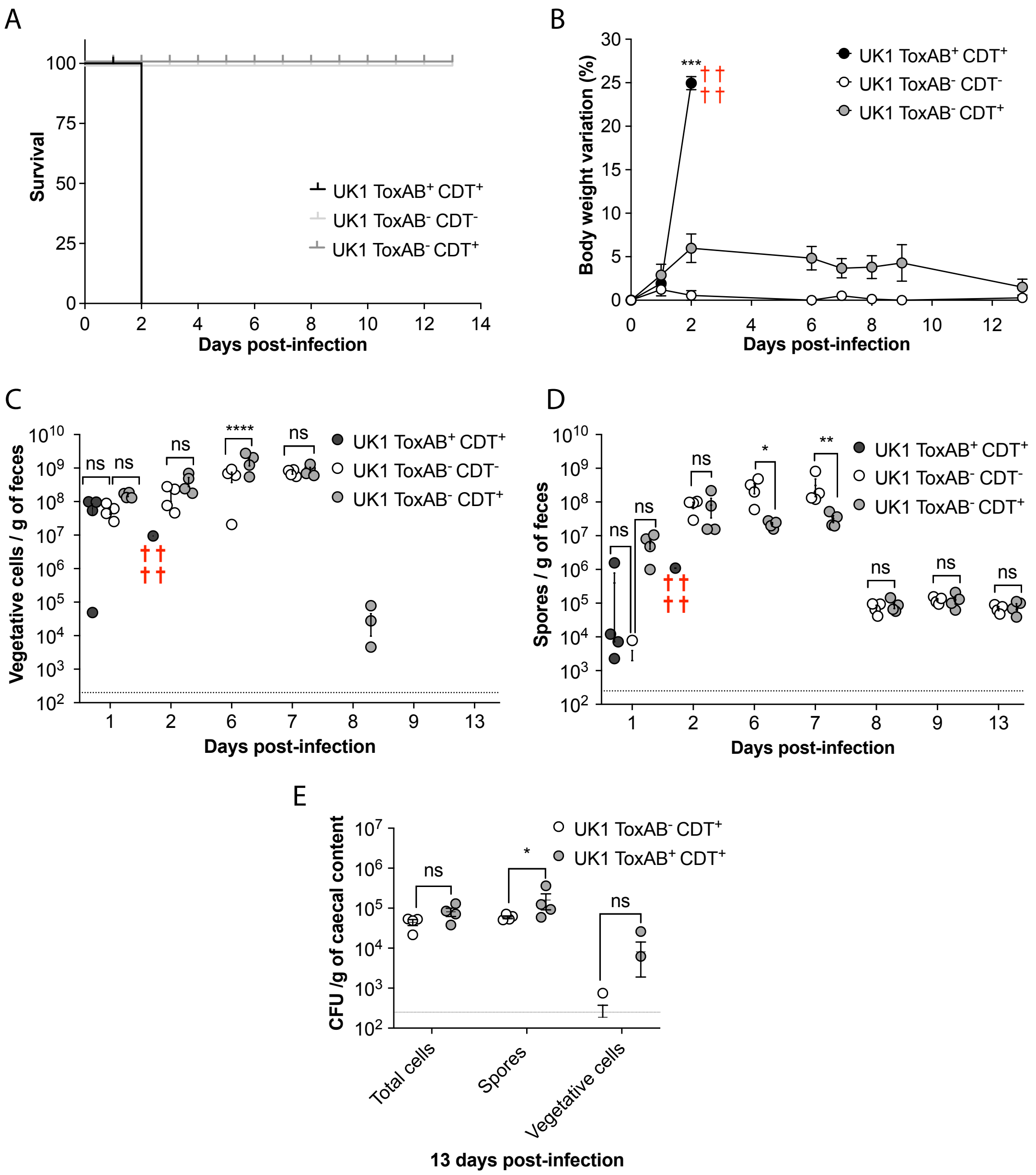


**Figure S7. Mice survival, body weight and *C. difficile* shedding into feces or caecal content.** C57Bl/6J Germ free mice 7-week old were infected with UK1 ToxAB$^+$CDT$^+$, UK1 ToxAB$^-$CDT$^-$ or UK1 ToxAB$^-$CDT$^+$. (A) Mice survival after infection with *C. difficile* strains as indicated. (B) Body weight average variation (%) of mice infected with *C. difficile* strains as indicated. (C) Total vegetative cells were analyzed different days p.i. (D) Spores were analyzed different days p.i. (E) Total cells, spores and vegetative cells detected from caecal content from mice infected with *C. difficile* strains as indicated 13 days p.i. Data represents mean with SEM. Multiple unpaired *t* tests were performed and statistical significance is represented with * p<0.05, ** p <0.01, ***p <0.001 and, **** p<0.0001. ns: no statistical significance.
